# Fc-engineered large molecules targeting blood-brain barrier transferrin receptor and CD98hc have distinct central nervous system and peripheral biodistribution compared to standard antibodies

**DOI:** 10.1101/2024.07.11.602993

**Authors:** Nathalie Khoury, Michelle E Pizzo, Claire B Discenza, David Joy, David Tatarakis, Mihail Ivilinov Todorov, Moritz Negwer, Connie Ha, Gabrielly L De Melo, Lily Sarrafha, Matthew J Simon, Darren Chan, Roni Chau, Kylie Chew, Johann Chow, Allisa Clemens, Yaneth Robles-Colmenares, Jason C Dugas, Joseph Duque, Doris Kaltenecker, Holly Kane, Amy Leung, Edwin Lozano, Arash Moshkforoush, Elysia Roche, Thomas Sandmann, Mabel Tong, Kaitlin Xa, Yinhan Zhou, Joseph W Lewcock, Ali Ertürk, Robert G Thorne, Meredith E K Calvert, Y Joy Yu Zuchero

## Abstract

The blood-brain barrier (BBB) poses a significant challenge drug delivery to the brain. BBB-crossing molecules are emerging as a new class of therapeutics with significant potential for central nervous system (CNS) indications. In particular, transferrin receptor (TfR)- and CD98 heavy chain (CD98hc)-targeting molecules have been demonstrated to cross the BBB for enhanced brain delivery. Previously, we reported TfR and CD98hc antibody transport vehicles (ATV^TfR^ and ATV^CD98hc^) that utilize these BBB receptors to improve CNS drug delivery^1,2^. Here, we provide a comprehensive and unbiased biodistribution characterization of ATV^TfR^ and ATV^CD98hc^ compared to a standard IgG at a multiscale level, ranging from whole-body to brain region- and cell type-targeting specificity. Mouse whole-body tissue clearing revealed distinct organ localization for each molecule. In the CNS, ATV^TfR^ and ATV^CD98hc^ not only achieves enhanced brain delivery but importantly, much broader parenchymal distribution in contrast to the severely limited distribution observed with a standard antibody that was not able to be improved even at very high dose levels. Using cell sorting and single-cell RNA sequencing of mouse brain, we revealed that standard IgG predominantly localizes to perivascular and leptomeningeal cells and reaches the CNS by entering the CSF, rather than crossing the BBB. In contrast, ATV^TfR^ and ATV^CD98hc^ enables broad parenchymal cell-specific distribution via transcytosis through brain endothelial cells (BECs) along the neurovasculature. Finally, we extended the translational relevance of our findings by revealing enhanced and broad brain and spinal cord biodistribution of ATV^TfR^ compared to standard IgG in cynomolgus monkey. Taken together, this multiscale analysis reveals in-depth biodistribution differences between ATV^TfR^, ATV^CD98hc^, and standard IgG. These results may better inform platform selection for specific therapeutic targets of interest, optimally matching platforms to desired CNS target engagement, peripheral organ exposures, and predict or potentially reduce off-target effects.

## Introduction

Disorders of the central nervous system (CNS) comprise a large area of unmet medical need. A major limitation to the success of many CNS treatments is the high selectivity and restrictiveness of the blood-brain barrier (BBB), which severely limits the effective delivery of antibodies and other large molecules into the brain ^3–8^. Despite numerous clinical trials for neurodegenerative diseases using large molecule therapeutics over the past decades, only a few recent efforts have shown clinical efficacy ^9–11^. Given that only around 0.01-0.1% of systemically introduced antibodies reach the brain, there is a great need to further improve the target engagement potential of CNS therapeutics by enhancing their brain uptake capacity and biodistribution ^1,12^.

The entry route of systemically delivered protein therapeutics into the brain remains incompletely understood. Brain endothelial cells (BECs) possess many unique cellular properties, including limited vesicular trafficking and preferential lysosomal degradation, both of which leads to very low levels of immunoglobulin (IgG) transport across BECs ^12,13^. Initial studies examining CNS biodistribution described endogenous serum proteins as distributing primarily to the leptomeninges and perivascular spaces in addition to the circumventricular organs ^14–16^, yielding distribution patterns similar to that resulting from exogenous protein administration into the CSF ^17^. Subsequent work has demonstrated a clear molecular size-dependent access of circulating proteins to the CSF under normal conditions, a finding historically interpreted as consistent with serum proteins primarily accessing the CSF across choroid plexus epithelial cells and nonchoroidal sites such as the circumventricular organs ^18^. Taken together, this suggests systemic and even direct CSF delivery of large molecules may often result in limited brain parenchymal distribution ^17^. Approaches for large molecule CNS delivery have sought to target highly expressed luminal proteins on BECs to enable receptor-mediated transcytosis or potentially other trafficking pathways. The most well-characterized of these BBB proteins are the transferrin receptor (TfR1 or TfR) and the more recently described CD98 heavy chain (CD98hc, also known as 4F2) ^1,2,19–27^. There has been an increase in the development of BBB-targeted large molecule platforms ^28–39^ as well as approvals of standard non-BBB targeted CNS antibody therapeutics^9–11^ yet an unbiased and comprehensive characterization of the precise pathways and cell type these BBB-targeting molecules distribute to in the CNS has not yet been explored. Additionally, a detailed biodistribution comparison of BBB-enabled molecules to traditional IgG has been lacking; it remains unclear whether the parenchymal biodistribution limitations of standard antibodies may be overcome simply with higher doses, or if their limited biodistribution is an inherent consequence of their specific route of entry into the brain.

We have previously described two BBB-crossing transport vehicles (TVs) that facilitate increased brain uptake and biodistribution by directly engineering the huIgG Fc to bind to either TfR or CD98hc (TV^TfR^, TV^CD98hc^) ^1,2^. The TV platform is highly modular, with applications for antibodies (ATVs), enzymes (ETVs), other proteins (PTVs), and oligonucleotide conjugates (OTVs) ^1,2,28,33–35,35^. The TV platform is engineered for distinct advantages compared to other brain delivery platforms, e.g. no unnatural linkers or appended sequences, retention of the native IgG architecture, and optimized TV affinity to maximize exposure ^1,2^. Several different TV fusion proteins have demonstrated efficacy in preclinical animal models of disease ^1,2,28,33–35,35^ and ongoing evaluation in clinical trials are on-going^40^. Although these prior preclinical reports have consistently described significant increases in brain exposure with TfR- and CD98hc-targeted TV molecules compared to standard IgG, a more comprehensive description of whole-body distribution and cell-type specific distribution of TVs throughout the brain has yet to be reported. Given that TfR and CD98hc are also expressed across peripheral organs, a more granular multilevel characterization is particularly important for optimized TV platform selection for various types of therapeutic targets. This knowledge would help to broaden the window between efficacy and safety, as well as to identify potential for targeting peripheral organs. Finally, more information is needed to better appreciate how distribution patterns and mechanisms revealed in rodent studies translate to primates.

Here, we address several of these questions using multiple novel technologies to characterize ATV and standard IgG biodistribution at a multiscale level from whole body to brain cellular subtypes. Whole-body tissue clearing identified organ-specific uptake of ATVs that was previously not appreciated, as well as enhanced ATV brain and spinal cord biodistribution to both vasculature and parenchyma. In contrast, control IgG predominantly localized to CSF-associated compartments, even at substantially higher doses, suggesting TV-enabled distribution via the vasculature is superior for widespread brain delivery. Brain cell-specific distribution using fluorescence activated cell sorting (FACS) and single cell RNA sequencing (scRNA-seq) demonstrate that ATV^TfR^ and ATV^CD98hc^ can be found in BECs and pericytes as they traffic through the BBB, with unique and consistent uptake in most parenchymal cell types. In contrast, control IgG predominantly distributed to CSF-bordering and perivascular cells associated with cerebral blood vessels, with minimal localization to BECs or parenchymal cell types. Importantly, we demonstrate that the vascular and parenchymal biodistribution of TV^TfR^-enabled molecules obtained in our mouse studies translated well to larger non-human primate brain and spinal cord.

## Results

### ATV^TfR^ and ATV^CD98hc^ exhibit distinct peripheral biodistribution patterns

Previous reports of peripheral distribution of TV-enabled molecules have provided useful insights, yet most approaches utilized to date have been focused upon preselected organs without spatial biodistribution information ^2,34^. We used whole-body tissue clearing and light sheet fluorescence microscopy (LSFM) to enable a more comprehensive and unbiased understanding of peripheral biodistribution ^41–43^. To isolate the contribution of TfR and CD98hc binding alone on brain exposure and biodistribution, ATV^TfR^ and ATV^CD98hc^ were engineered with non-targeting Fabs as well as with Fc mutations to eliminate binding to Fc gamma receptor and complement proteins (L234A/L235A/P329G) ^44^ (**Fig 1a)**. Control IgG, ATV^TfR^, or ATV^CD98hc^ were pre-conjugated with Alexa Fluor 647 (AF647) and systemically administered into wild-type, TfR^mu/hu^ KI, or CD98hc^mu/hu^ KI mice followed by whole body tissue clearing and 3D LSFM (**Fig 1b)**. Terminal time points were selected based on previously reported brain Cmax for each platform ^1,2^. Dorsal and ventral 3D views, as well as rotating and sectional fly-through video, revealed unique biodistribution profiles for each molecule (**Fig 1c, Video 1**). Quantification of the AF647 signal from ATV^TfR^ and ATV^CD98hc^ (normalized to control IgG) revealed that both ATVs are taken up at higher levels across numerous peripheral organs with differentiated distribution patterns (**Fig 1d-e**). ATV^TfR^ preferentially localized to bones (e.g. tibia, shoulder bone, vertebra, and skull), consistent with high TfR expression in erythroid precursor cells within the bone marrow (**Fig 1d-f, Video 1)**, ^45,46^. The brain exhibited the second highest level of ATV^TfR^ uptake, with some slight regional differences observed. Additional organs that showed pronounced uptake of ATV^TfR^ included the large intestine, kidney (cortex and medulla), lacrimal glands, lung, and spleen. Intriguingly, the organ that showed the highest level of ATV^CD98hc^ uptake was the lacrimal gland which, in addition to the parotid gland, and sciatic nerve, have not been previously identified as peripheral targets for CD98hc-binding constructs (**Fig 1e, g, Video 1**). Consistent with previous reports, enhanced uptake of ATV^CD98hc^ was also observed in the large intestine, kidneys, pancreas, and testes, in addition to the brain ^2^. As with ATV^TfR^, the brain exhibited widespread and significant uptake of ATV^CD98hc2^. These peripheral distribution data suggest that TVs may not only serve as brain delivery platforms but could potentially also be exploited in certain cases to provide enhanced delivery to specific peripheral organs.

**Figure 1.**
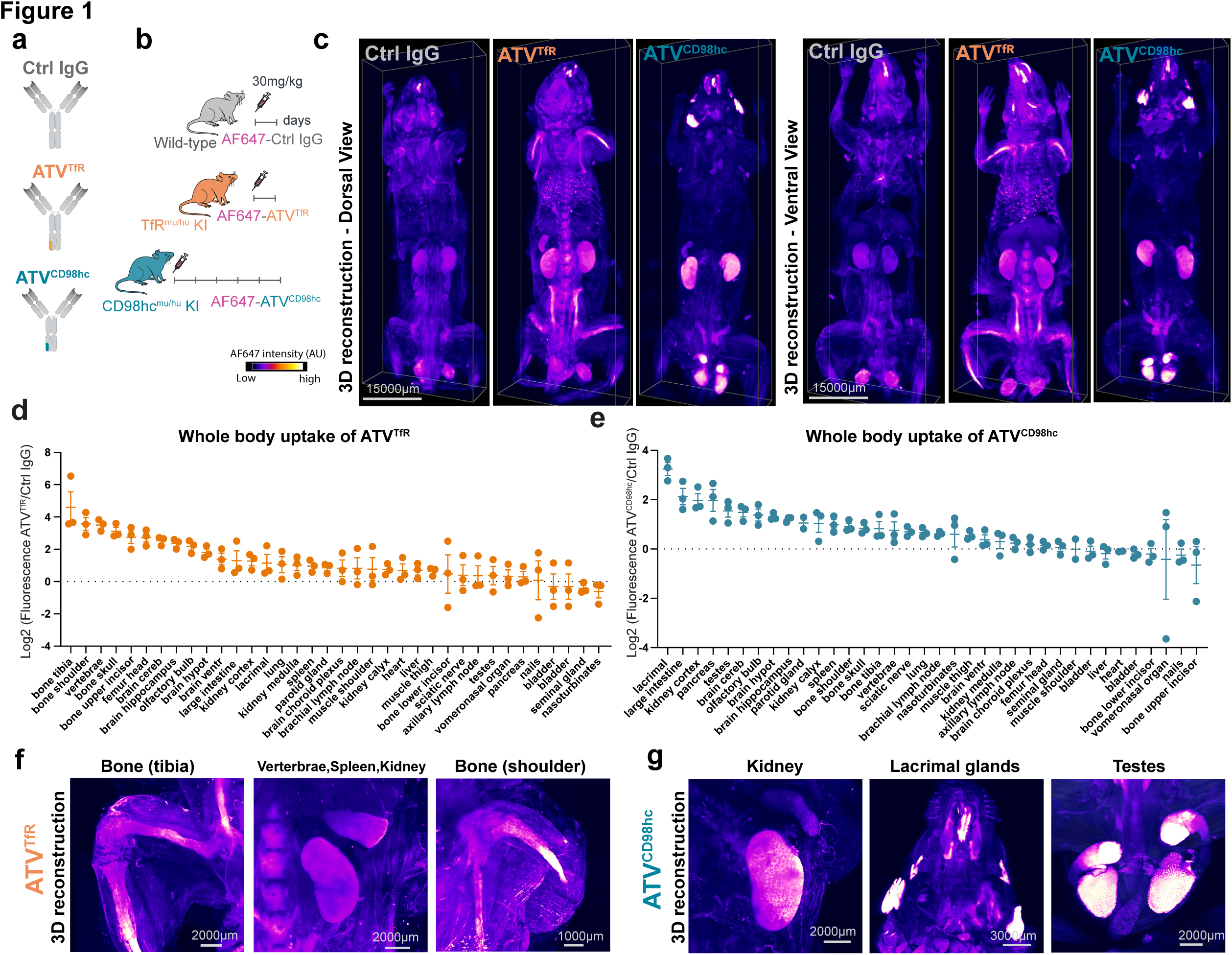
Whole body mouse fluorescence imaging reveals unique peripheral biodistribution patterns of ATV^TfR^, ATV^CD98hc^, and control IgG. **a.** Schematic of the molecules used with non-targeting control Fabs. The orange patch in ATV^TfR^ Fc region binds to TfR and blue patch in ATV^CD98hc^ binds to CD98hc. **b.** Schematic of the experimental paradigm. TfR^mu/hu^ KI, CD98hc^mu/hu^ KI, or WT mice were dosed with 30 mg/kg AF647-conjugated control IgG and ATV^TfR^ for 1 day and ATV^CD98hc^ for 5 days. **c.** Ventral and dorsal view 3D immunofluorescence images AF647-conjugated control IgG, ATV^TfR^, or ATV^CD98hc^ in the whole mouse. Representative immunostaining from n = 3/group. **d-e.** Quantification of fluorescence intensity of AF647-ATV^TfR^ **(d)** or AF647-ATV^CD98hc^ **(e)** normalized to AF647-conjugated control IgG. n = 3/ group, mean ± sem. **f-g.** 3D reconstructed images of select peripheral organs with high ATV^TfR^ **(f)** or ATV^CD98hc^ **(g)** uptake. Representative images from n = 3/group.

### 3D imaging reveals enhanced brain biodistribution of ATV^TfR^ and ATV^CD98hc^

Whole-body tissue clearing also provided the opportunity to further evaluate the biodistribution of ATV^TfR^ and ATV^CD98hc^ compared to standard IgG throughout the brain, meninges, and other associated tissues in the intact mouse head. 3D reconstructed images revealed striking differences in the overall distribution of ATVs compared to control IgG. The latter exhibited a clear signal primarily at the brain surface, corresponding to leptomeningeal tissue and associated blood vessels on the brain surface (**Fig 2a**). Weaker IgG signal was also evident in the lateral ventricles. In contrast, the ATV^TfR^ and ATV^CD98hc^ groups exhibited strong and distinct patterns of signal intensities throughout the brain (**Fig 2a, Videos 2-4**). Quantification of the AF647 signal from the brain surface extending through deeper cortical regions revealed a consistently elevated signal for both ATVs, in stark contrast to the low level of signal for control IgG (**Fig 2b**).

**Figure 2.**
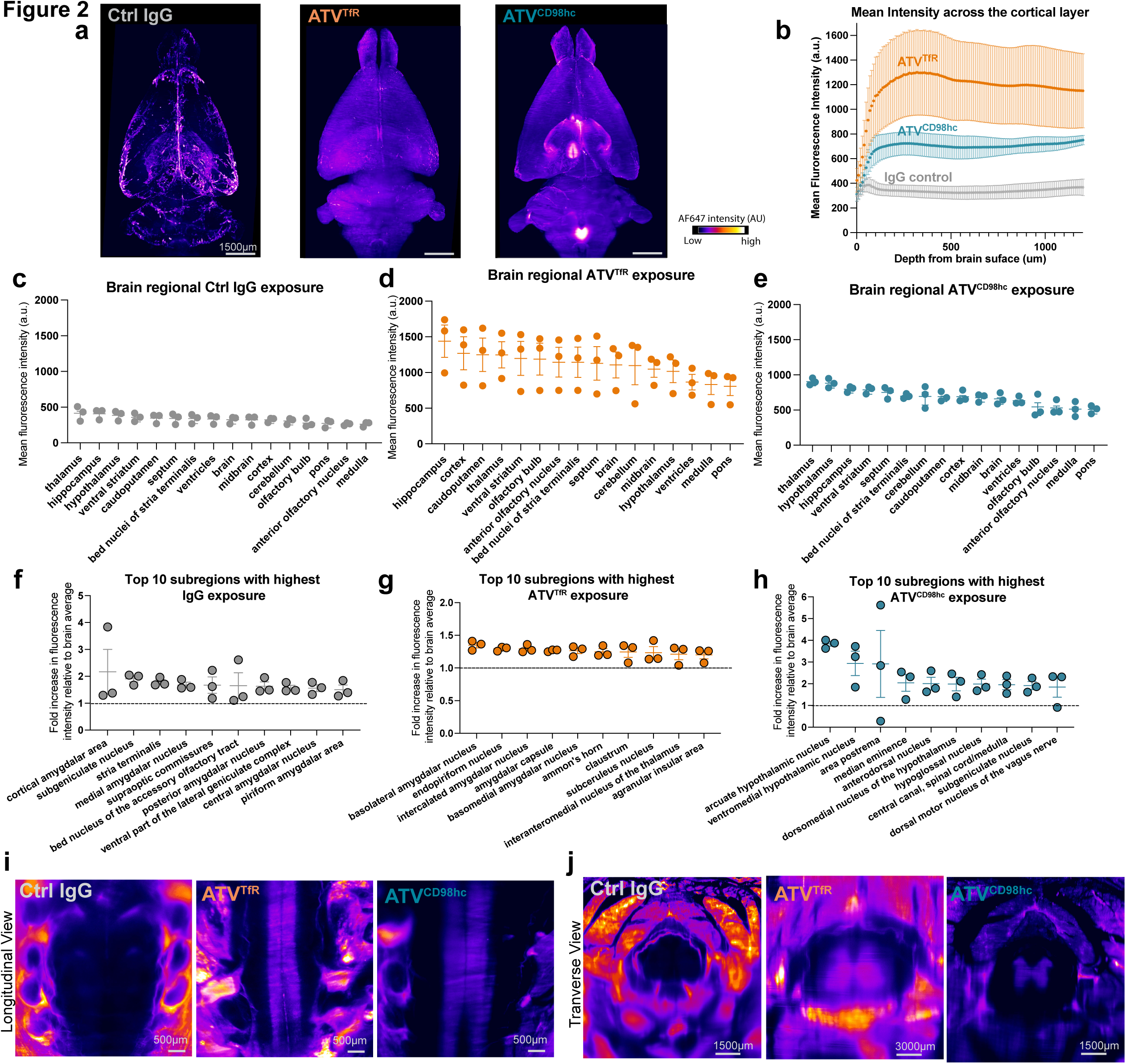
Enhanced brain uptake and biodistribution of ATV^TfR^ and ATV^CD98hc^ in the mouse brain and spinal cord by the whole-body fluorescence imaging. **a.** Representative images from AI-segmented 3D-reconstructed brains obtained from the whole-body tissue cleared mice dosed with AF647-conjugated control IgG, ATV^TfR^, or ATV^CD98hc^. Representative images from n = 3/group **b.** Mean fluorescence intensity (a.u.) of control IgG, ATV^TfR^, and ATV^CD98hc^ as a function of brain depth from the cortical surface (0 μm) moving into deeper cortical tissues (1,200 μm). n = 3/ group, mean ± sem. **c-e.** Mean fluorescence intensity (a.u.) of AF647-conjugated control IgG **(c)**, ATV^TfR^ **(d**), or ATV^CD98hc^ **(e)** across different mouse brain regions from the whole-body tissue cleared mice. n = 3/ group, mean ± sem. **f-h.** Top ten regions with highest fold change in fluorescence intensity compared to brain average fluorescence intensity for AF647-conjugated IgG **(f)**, ATV^TfR^ **(g**), or ATV^CD98hc^ **(h)**. Dotted line denotes the average brain intensity. n = 3/ group, mean ± sem. **i-j.** Optical slice longitudinal **(i)** or transverse (**j**) views of fluorescence images from mouse spinal cord from whole-body tissue cleared mice. Representative images from n = 3/group.

As many neurological disorders affect distinct brain regions, we next determined whether the two ATV platforms exhibited differential uptake across brain regions that could be informative for targeting of specific therapeutics. As expected, control IgG administration yielded a consistently low signal across all AI (artificial intelligence)-segmented brain regions, in marked contrast to the higher intensities across all regions associated with both ATV molecules **(Fig 2c-e)**. A global assessment revealed a relatively homogenous exposure for ATV^TfR^ across the whole brain, albeit with slightly more enhanced uptake in the hippocampus and cortex **(Fig 2d)**. ATV^CD98hc^ also exhibited a relatively homogenous exposure and revealed regions with elevated exposure such as the thalamus, hypothalamus, and hippocampus **(Fig 2e)**. Since this initial quantification approach covered large brain areas, an unbiased approach identified additional smaller subregions with the highest exposure levels relative to the average whole brain signal **(Fig 2f-h)**. Subregion-specific enhancement was found to be modest for control IgG and ATV^TfR^. In contrast, ATV^CD98hc^ distribution revealed several subregions with IgG concentrations more than two-fold above the average signal in brain, including the arcuate hypothalamic nucleus, the ventromedial hypothalamic nucleus, area postrema, and median eminence, among others **(Fig 2e).** The high uptake of ATV^CD98hc^ observed in these circumventricular organs (arcuate hypothalamic nucleus, ventromedial hypothalamic nucleus, area postrema, and median eminence) is consistent with high expression in these regions of CD98hc and one of its binding partners, LAT1 (L-type amino acid transporter) ^47–49^.

ATV uptake was also assessed in the spinal cord, a region particularly affected in CNS diseases such as amyotrophic lateral sclerosis, multiple sclerosis, and spinal muscular atrophy, among others. Both longitudinal (**Fig 2i**) and cross-sectional (**Fig 2j**) views of the spinal cord showed clear and enhanced exposure of both ATV^TfR^ and ATV^CD98hc^ compared to control IgG, which did not show detectable signal. Additionally, the cross-sectional images revealed that within the ATV groups, uptake was mostly evident in the spinal gray matter (**Fig 2j**). Taken together, these findings further support the higher exposure and distribution of ATV^TfR^ and ATV^CD98hc^ across CNS regions.

### ATV^TfR^ and ATV^CD98hc^ exhibit enhanced brain exposure and biodistribution

Although higher total brain concentration can be achieved with standard IgG through using very high systemic doses, it remains unclear whether dosing higher can result in a homogeneous brain biodistribution achieved with ATVs. To test this, we first compared brain exposure of control IgG, ATV^TfR^, or ATV^CD98hc^ that were pre-conjugated with AF647 and systemically administered into wild-type, TfR^mu/hu^ KI, or CD98hc^mu/hu^ KI mice, respectively (**Fig 3a)**. Bulk brain exposure levels for ATV^TfR^ and ATV^CD98hc^ were significantly higher compared to control IgG when each molecule was administered at the same dose (**Fig 3b**). Consistent with our whole-body tissue clearing, widefield fluorescence imaging of whole sagittal brain sections by IHC revealed widespread vascular and parenchymal localization of ATV^TfR^ and ATV^CD98hc^ across the whole brain, while control IgG was primarily localized to the choroid plexus and leptomeningeal tissues at the brain surfaces (**Fig 3c**). To investigate whether increasing the dose could overcome this limited control IgG brain distribution, we dosed either 100 mg/kg of control IgG or 15 mg/kg of ATV^TfR^, which resulted in comparable bulk brain concentrations (**Fig 3d**). However, the biodistribution pattern of both molecules remained similar to the dose-matched fluorophore-conjugated molecules (**Fig 3 c,e**). That is, control IgG exposure remained predominantly limited to the pial brain surface and appeared remarkably similar to what has been reported when IgG is applied directly into the CSF by intracisternal administration ^17^. High signal was also observed in association with the choroid plexus, particularly in the stroma on the blood-side (basolateral aspect) of the polarized epithelial cells (**Supplementary** Fig 1a). Putative perivascular signal was observed, particularly prominent in the colliculi, hindbrain, and the thalamus adjacent to the lateral ventricle (**Fig 3f**). We observed a prominent signal in the fimbria of the hippocampus bordering the ventricle and a gradient of diffuse parenchymal signal in the thalamus which appeared to arise from the lateral ventricle (**Fig 3f**). Control IgG signal was also prominently observed around the olfactory nerve layer of the ventral olfactory bulb, near a known drainage pathway for CSF (**Fig 3e, Supplementary** Fig 1b) ^17,50–52^. Despite the high dose of control IgG, the parenchyma appeared to have little signal relative to ATV^TfR^. Notably, ATV^TfR^ localized to capillary and venular endothelial cells while mostly absent from arterial endothelial cells (e.g. notably darker putative arteriole based on morphology in neocortex), a pattern that aligns with vascular zonation differences in TfR expression (**Fig 3f**) ^53^. ATV^TfR^ signal was also observed in the choroid plexus, but with a more diffuse pattern and relatively lower intensity compared to control IgG.

**Figure 3.**
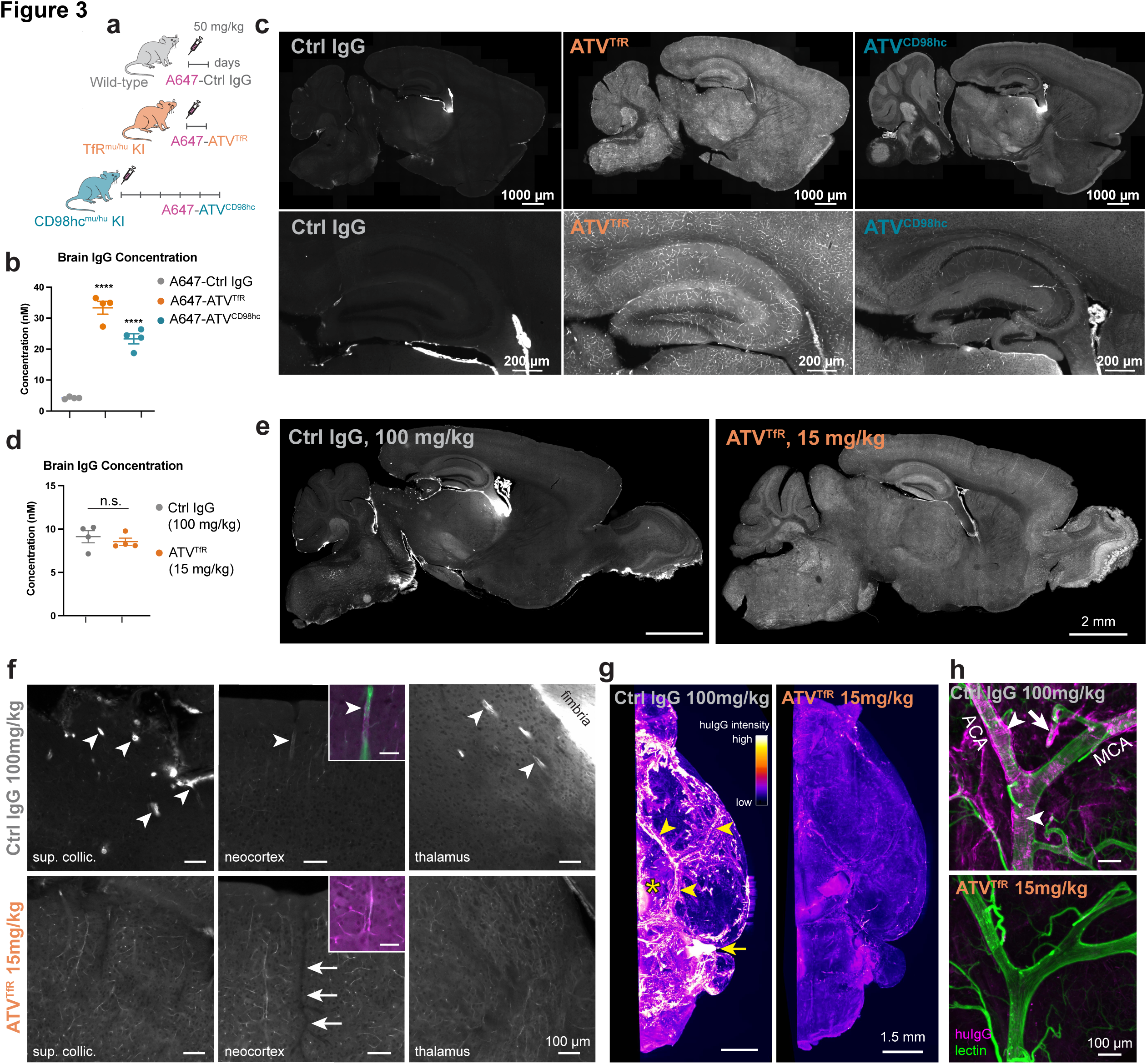
ATV^TfR^ and ATV^CD98hc^ exhibit enhanced brain uptake and biodistribution compared to control IgG. **a.** Schematic of the experimental design. WT, TfR ^mu/hu^ KI, and CD98hc^mu/hu^ KI mice were dosed with 50 mg/kg of AF647-conjugated control IgG and ATV^TfR^ for 1 day and ATV^CD98hc^ for 5 days. **b.** Brain concentration of AF647-conjugated control IgG, ATV^TfR^, or ATV^CD98hc^ as measured by huIgG ELISA in bulk brain lysates after a single 45 mg/kg IV dose. n = 4/ group, mean ± SEM, one-way ANOVA ****p < 0.0001 compared to control IgG. **c.** Representative immunofluorescence in sagittal brain sections of AF647-conjugated control IgG, ATV^TfR^, and ATV^CD98hc^ in mouse. Representative immunostainings from n = 2/group. **d.** Brain lysate concentration measured by ELISA one day after a single 100 mg/kg IV dose of control IgG or 15 mg/kg of ATV^TfR^ (n=4 mice/group). **e.** Immunodetection of huIgG in whole sagittal brain sections by widefield imaging. **f.** Magnified examples of brain regions from (e) including the superior colliculus, neocortex, and thalamus. Examples of putative perivascular huIgG signal indicated by arrowheads; notably darker putative arteriole indicated by arrows. Insets show immunodetected huIgG (magenta) and vascular marker caveolin-1 (green), scalebar 50 μm. **g**. Ventral volume view of immunodetected huIgG in tissue-cleared whole mouse hemibrain (n=6 mice/group). Arrowheads indicate huIgG associated with the large surface arteries, asterisk indicates the median eminence, arrow indicates cranial nerves (region of trigeminal, facial, and vestibulocochlear nerves). **h.** Higher magnification light sheet imaging of immunodetected huIgG and vascular marker lectin around the middle cerebral artery (MCA) and anterior cerebral artery (ACA) branching from the circle of Willis. Circumferential banding pattern around putative smooth muscle cells indicated by arrowheads; arrow indicates putative perivascular profile of a penetrating vessel. For micrographs, display settings were optimized independently for each region of interest and were identical for both treatment groups.

We next took advantage of the prominent signal achieved with the high 100 mg/kg dose of control IgG to better visualize distribution within the leptomeninges and perivascular spaces of surface vessels in 3D by brain tissue clearing and LSFM imaging. These regions are often difficult to evaluate spatially by 2D IHC but may have particular safety implications for certain classes of CNS antibody therapeutics ^54–56^. Perfusion with lectin was used to enable visualization of the entire brain vasculature. Control IgG showed a clear and substantial association with the brain surface (presumably associated with leptomeningeal cells of the arachnoid and pia), leptomeningeal vessels (putative perivascular signal), the median eminence (a circumventricular organ lacking a blood-brain barrier), and hindbrain cranial nerves (putative routes for CSF clearance), while ATV^TfR^ did not strongly localize to any of these compartments (**Fig 3g**). Higher resolution imaging of the middle cerebral artery as it branches from the circle of Willis revealed control IgG localization on the abluminal (CNS) side of the vasculature **(Fig 3h)**. The signal appeared to be perivascular, with a circumferential banding pattern around smooth muscle cells that suggests control IgG could be accessing these spaces from the CSF. In contrast, ATV^TfR^ was minimally associated with these leptomeningeal and perivascular compartments. A lateral volume view of the middle cerebral artery and the underlying parenchyma showed a stark drop-off in control IgG signal within the parenchyma, suggesting inefficient penetration from the CSF and leptomeninges, aside from occasional perivascular signal evident around penetrating vessels (**Supplementary** Fig 1e-f).

### ATV^TfR^ and ATV^CD98hc^ localize to brain vascular and parenchymal cells while control IgG localizes to blood-CSF barrier and perivascular BBB cells

Obtaining unbiased brain cell-specific biodistribution patterns of ATV^TfR^, ATV^CD98hc^, and control IgG by IHC faces numerous challenges, particularly detection in less abundant cell types. We sought to address this by devising a highly sensitive and semiquantitative approach that allowed characterization of brain cellular biodistribution in an unbiased and comprehensive manner. Mice were dosed with AF647-conjugated ATV^TfR^, ATV^CD98hc^, or control IgG, and brains were dissociated and sorted by flow cytometry based on AF647 intensity. Terminal time points were selected based on previously reported brain Cmax for each platform (1 day for control IgG, ATV^TfR^, or 5 days for ATV^CD98hc 1,2^ )(**Fig. 4a**). Flow cytometry revealed that only around 1% of the live brain cells from the control IgG group were AF647-positive in contrast with substantially higher percentages for ATV^TfR^ and ATV^CD98hc^ (**Fig. 4b**). To generate semiquantitative distribution data, we divided the AF647-positive into low, medium, and high bins by dividing the total signal from the ATV^TfR^ group equally into thirds, such that each AF647-positive bin contained approximately 33% of the total AF647-positive cells (**Fig. 4b**). We then applied these gating cutoffs to the remaining treatment groups to enable comparisons across groups. Single cell RNA-seq was performed on all four bins (i.e. AF647-low, -medium, and -high bins, along with a negative bin denoting negligible signal) allowing us to identify and quantify AF647-labeled molecules within cell types that constitute each bin.

**Figure 4.**
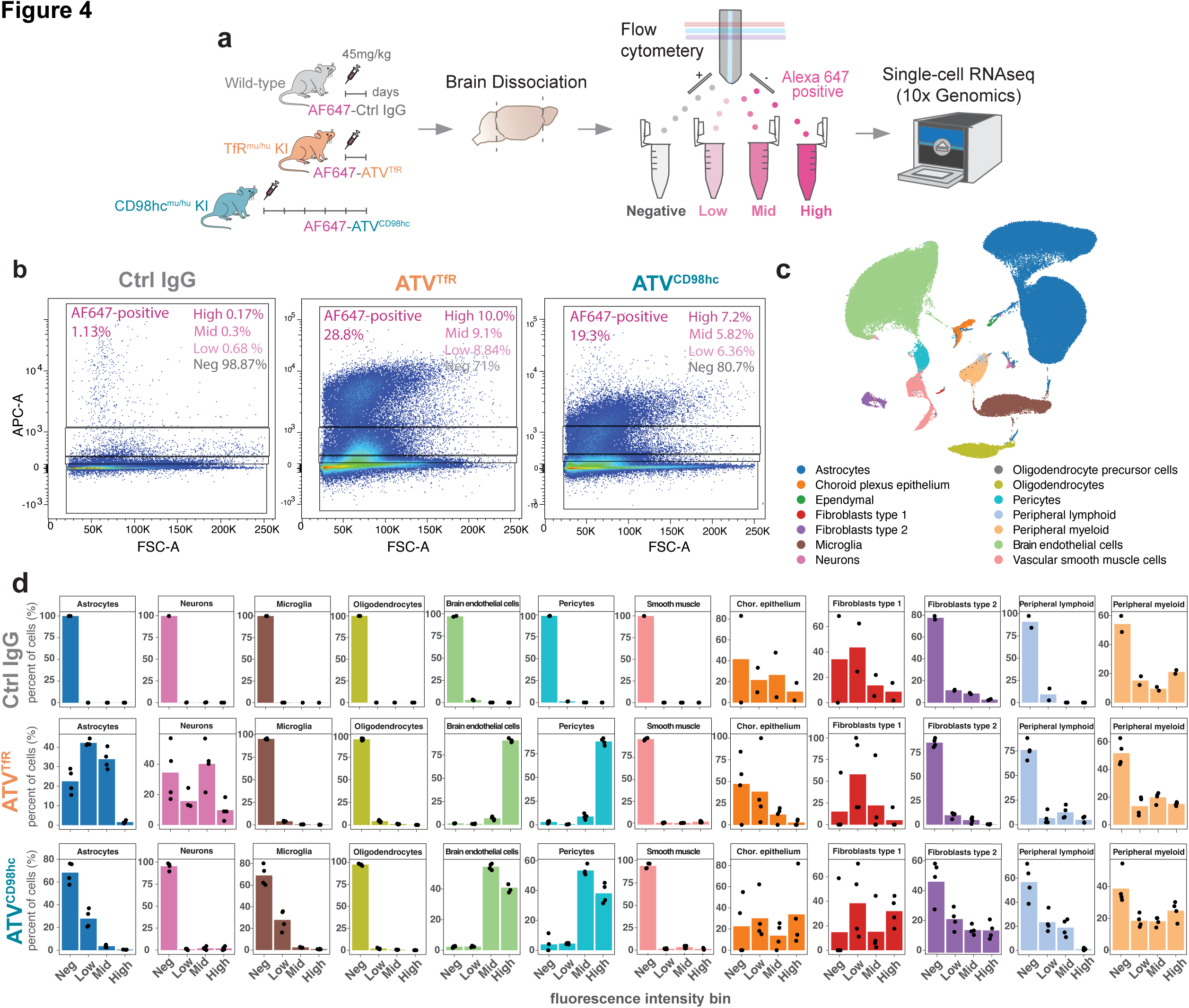
ATV^TfR^ and ATV^CD98hc^ localize to BBB vascular and CNS parenchymal cells while control IgG localizes to blood-CSF and perivascular barrier cells. **a.** Schematic of the experimental design: Single cells obtained from dissociated brains of WT, TfR ^mu/hu^ KI, and CD98hc^mu/hu^ KI mice dosed with 45 mg/kg AF647-conjugated antibodies were sorted based on AF647 signal and then loaded onto the 10x Genomics platform for scRNA sequencing. **b.** Representative flow cytometry plots showing the percentage of AF647-positive cells and their percent distribution across the negative, low, mid, and high AF647 intensity bins. **c.** UMAP of cell clusters captured from the dissociated brains of mice from all three treatment groups and across all four bins. **d.** Bar graphs depict mean percent of total huIgG distribution across indicated cell types captured per treatment group across fluorescence intensity bins. Points represent individual mice. Control IgG (n = 2/group); ATV^TfR^ and ATV^CD98hc^ (n = 4/ group).

Dimensionality reduction and predictive cell type labeling across all samples revealed that our approach captured not only major cell types of the brain such as neurons, microglia, astrocytes, and oligodendrocytes, but also cells of the neurovascular unit such as endothelial cells, pericytes, and vascular smooth muscle cells (VSMC), along with other border-associated cell types including perivascular lymphoid and myeloid cells, choroid plexus epithelial cells (CPECs) and two types of fibroblast cells (**Fig 4c; Supplementary** Fig. 2b). We divided fibroblasts into two broad types for ease of evaluation given the parameters of our dataset: type 1 fibroblasts predominantly expressed markers of the recently reported brain fibroblast (BFB) subtypes including dural border and arachnoid barrier fibroblasts (BFB5, BFB4 and BFB3), while type 2 fibroblasts expressed markers associated with the inner arachnoid, pia, perivascular compartments, and choroid plexus (BFB3, BFB2, BFB1b, BFB1a, and BFB6) ^57^ (**Supplementary** Fig. 2c). Notably, no control IgG was observed in any major parenchymal cell types including astrocytes, neurons, microglia, and oligodendrocytes, as these cell types were only found in the AF647-negative bin **(Fig 4d)**. The vast majority (97%) of vascular endothelial cells (i.e., BECs), pericytes, and vascular smooth muscle cells (VSMCs) also contained negligible levels of control IgG, suggesting minimal trafficking across the BBB. The only cell types where any appreciable control IgG was detected were the choroid plexus epithelial cells (inner blood-CSF epithelial barrier), type 1 and 2 fibroblasts associated with the meninges and perivascular compartments, and perivascular lymphoid and myeloid cells (**Fig 4d**).

ATV^TfR^ and ATV^CD98hc^ biodistribution exhibited a markedly different cellular biodistribution pattern. A significant percentage of ATV^TfR^ and ATV^CD98hc^ localized to both vascular and parenchymal cell types, where the vast majority of BECs and pericytes in both ATV^TfR^ and ATV^CD98hc^-treated groups fell in the AF647-mid and -high bins (**Fig 4d**). In addition, ATVs displayed localization to cells of the blood-CSF barriers (choroid plexus epithelial cells), fibroblasts and immune cell populations (**Fig 4d**). Each ATV exhibited unique parenchymal biodistribution characteristics as well. ATV^TfR^ exhibited strong localization to astrocytes and neurons, along with minimal but measurable localization to microglia and oligodendrocytes. Despite the very low cell numbers of neurons captured (279 cells in total across treatments), we were able to discern distinct localization patterns between ATV^TfR^ and ATV^CD98hc^ in this cell type, broadly consistent with previous reports by IHC ^1,2^. Specifically, higher localization of ATV^TfR^ was found in neurons, whereas ATV^CD98hc^ was rarely detected in this population (**Fig 4d**) but instead exhibited appreciably more localization to microglia compared to ATV^TfR^. One caveat of our approach was an expected loss of astrocytic processes due to the single-cell dissociation protocol. As we previously observed high expression of CD98hc on aquaporin 4-positive astrocytic endfeet and processes^2^, it is likely that our current approach underestimates localization of ATV^CD98hc^ to astrocytes. Taken together, our application of a novel method for evaluation of brain cell biodistribution revealed a highly specific cell-type distribution pattern of each ATV, compared to the limited parenchymal cell distribution of control IgG.

### ATV^TfR^ and ATV^CD98hc^ exhibit localization to BECs across the arterio-venous spectrum

Distinct expression patterns of transporters along the arterio-venous axis have been previously described in both mice and humans ^53,58^. To determine if ATVs are differentially taken up by BEC subtypes, we next subclustered BECs into arterial, capillary, and venous populations based on markers established in previous mouse single-cell studies ^59,60^ (**Supplementary** Fig. 2d-e) and determined the distribution of molecules to these subtypes across the intensity bins (**Fig 5**). There was minimal localization of control IgG to BECs, and among the ∼2% found in the AF647-low bin, nearly all signal belonged to the venous population (**Fig 5a).** Consistent with active trafficking across the BBB, ATV^TfR^ localized strongly to capillary and venous cells, with greater than 80% of AF647-positive cells falling within the high bin for these BEC subtypes, and a lower proportion (∼60%) falling within the high bin for arterial cells (**Fig 5b**). ATV^CD98hc^ exhibited a more similar distribution across all three BEC subtypes, with ∼50% of AF647-positive cells falling within both mid and high bins (**Fig 5c**). Overall, these data are largely consistent with their expression along the arterio-venous axis as previously reported in both mice and human transcriptomics datasets, and suggests ATV^TfR^ and ATV^CD98hc^ are actively trafficked across the BBB multiple through multiple BEC subtypes ^53,58^.

**Figure 5.**
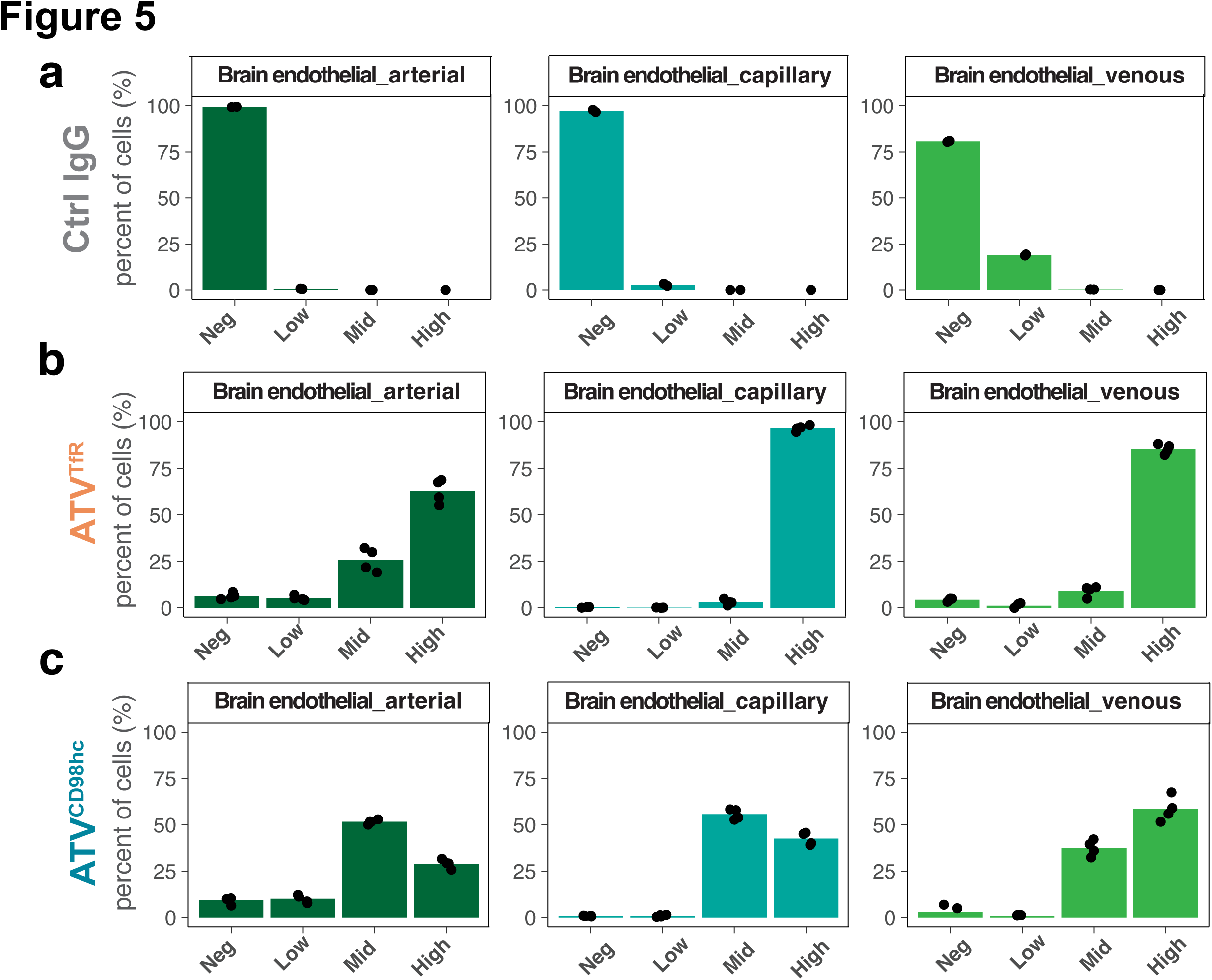
ATV^TfR^ and ATV^CD98hc^ exhibit distinctly different localization to BEC subtypes compared to control IgG across the arteriovenous axis. **a-c** Bar graph depict mean percent of total huIgG distribution across arterial, capillary, and venous endothelial cells captured per treatment group across fluorescence intensity bins. Points represent individual mice. Control IgG (n = 2/group); ATV^TfR^ and ATV^CD98hc^ (n = 4/ group).

### Enhanced vascular and parenchymal biodistribution patterns of ATV^TfR^ in cynomolgus monkey brain and spinal cord

Our previous work established significant increases in bulk brain exposure ATV in non-human primates, though a more granular biodistribution analysis across the entire brain and within more discrete brain regions has yet to be evaluated ^1^. Given TfR-based approaches are the most clinically advanced and widely used brain delivery platforms, we sought to examine whole brain biodistribution of ATV^TfR^ versus control IgG using tissue clearing and 3D LSFM imaging on the brains and spinal cords of cynomolgus monkeys. Animals were dosed systemically with AF647-conjugated control IgG or ATV^TfR^ and brain vasculature were simultaneously labeled using wheat germ agglutinin (**Fig 6a**)^61^. Given the technical complexity of the experiment and desire to minimize NHP lives, a single monkey per treatment was used. A global 3D view of the hemibrain revealed higher signal for ATV^TfR^ throughout the entire brain compared to control IgG (**Fig 6b, c**). Coronal slice views and higher magnification images further revealed clear vascular and parenchymal signals for ATV^TfR^ within the smallest microvessels (i.e., capillaries), while control IgG was primarily localized only to the perivascular space of larger penetrating vessels (**Fig 6d-g; Supplementary** Figure 3a-b). This biodistribution pattern was evident across multiple brain regions (**Supplementary** Figure 3aj**-d**), in coronal sections throughout the whole brain (**Supplementary** Figure 4), and sectional fly-through 3D videos (**Videos 5-6**). In all areas and regions of interest (ROIs) examined, delivery of ATV^TfR^ to the cynomolgus monkey brain clearly exceeded that of control IgG. The mean fluorescence of ATV^TfR^ was higher than control IgG in both segmented brain vessels as well as in the non-vessel parenchymal fraction (**Fig 6h-i**). Indeed, the ATV^TfR^ distribution reached deep into the tissue as evidenced by the persistence of signal intensity in deeper brain regions (**Fig 6i**). Significantly higher ATV^TfR^ signal was found in smaller caliber vessels (<40 µm) compared to larger vessels (>60um), consistent with the predominant capillary and venule TfR BEC distribution patterns that have been reported in mice as well as our BEC subtype cell distribution (**Fig 6j**; **Fig. 5**) ^53,62^. ATV^TfR^ signal was higher than control IgG across all brain regions measured, with the largest differences observed in the cortex, diagonal subpallium, amygdala, hypothalamus, and pons (**Fig 6k**). Isolated cervical spinal cords also revealed striking levels of ATV^TfR^ uptake in both the vasculature and parenchyma, with notably higher signal in the spinal gray matter **(Fig 6m)**. In contrast, control IgG signal in the spinal cord was notably faint, with prominent signal primarily observed only in spinal nerve roots (**Fig 6l**), comprising another putative CSF drainage pathway and similar to what has been observed following intrathecal antibody administration ^17,50^. Taken together, these findings in cynomolgus monkey brain and spinal cord illustrate how ATV^TfR^ enables both higher CNS exposure as well as significantly broader and homogenous biodistribution throughout the parenchyma compared to non-BBB targeted IgG in larger primate brains.

**Figure 6.**
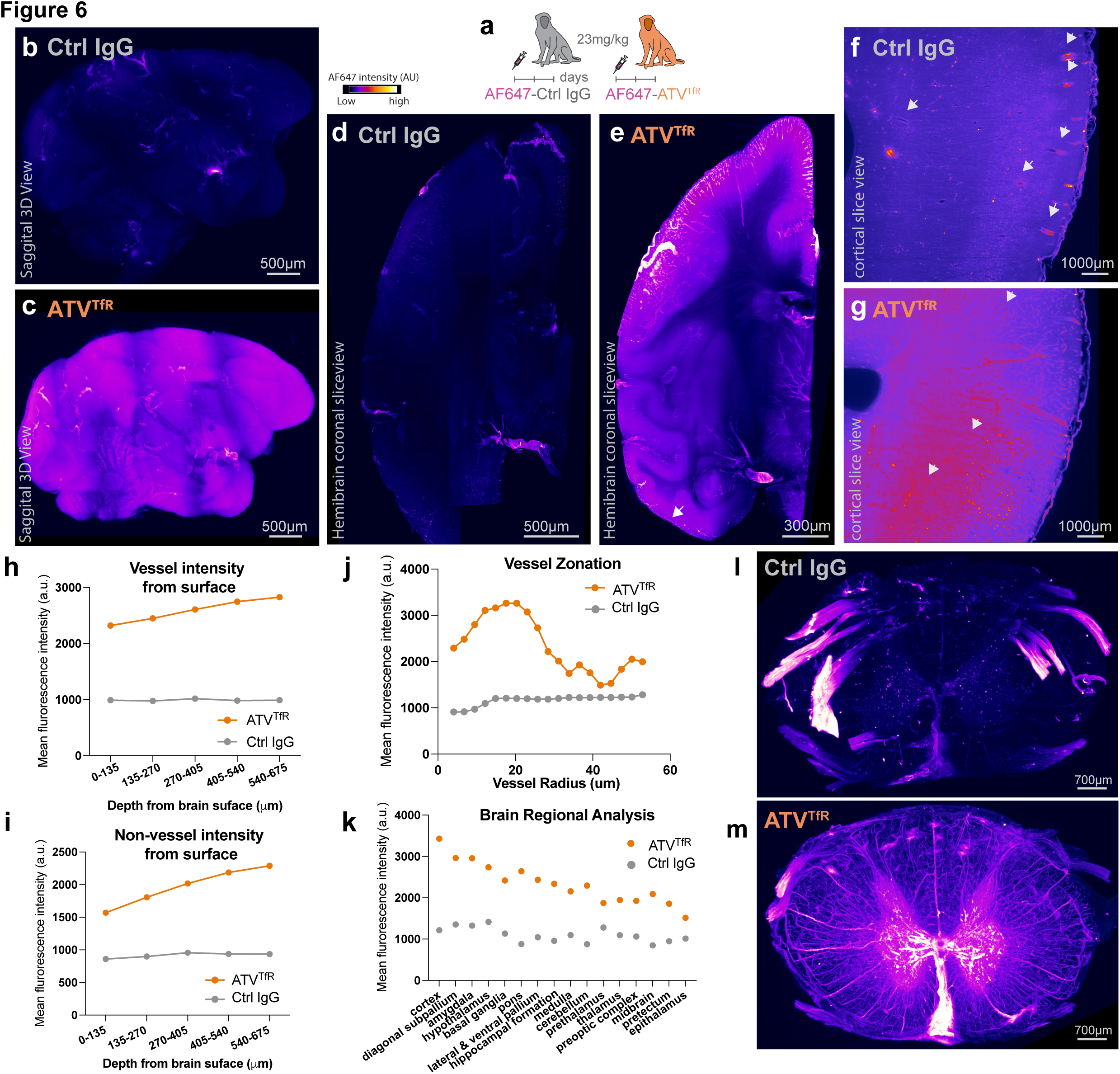
Enhanced exposure and biodistribution of ATV^TfR^ in cynomolgus monkey brain and spinal cord. **a.** Schematic of the *in vivo* experimental design. Immunofluorescence images of 3D reconstructed hemibrains (1x objective) **(b,c)**, coronal slice view of hemibrain (200μm thick, 1x objective) **(d,e)**, and cortical slice view of temporal lobe (ventral cortex) (200μm thick, 4x objective) **(f,g)** from cynomolgus monkeys dosed with 23 mg/kg AF647-conjugated control IgG or ATV^TfR^ (IV) for 2 days. Arrow heads in **(f**) highlight localization of control IgG to large penetrating vessels while arrow heads in **(g)** highlight localization of ATV^TfR^ within large vessels as well as capillaries. Immunostaining image from n = 1/group. **h-i.** Mean fluorescence intensity of ATV^TfR^ and control IgG in vessels **(h)** and non-vessels **(i)** in cynomolgus monkey brains across regions of different depths from surface of the brain. **j.** Mean fluorescence intensity of ATV^TfR^ and control IgG within vessels of different radii in the cynomolgus monkey brains. **k.** Mean fluorescence intensity of ATV^TfR^ and control IgG across different brain regions in cynomolgus monkeys. **l-m.** 3D Immunofluorescence images of spinal cords from cynomolgus monkeys dosed with AF647-conjugated control IgG **(l)** or ATV^TfR^ **(m).** Immunostaining image from n = 1/group.

## Discussion

BBB-crossing molecules are emerging as a new class of therapeutics with huge potential for CNS indications. In this study, we provide a comprehensive and unbiased examination of the biodistribution of standard antibody, ATV^TfR^, and ATV^CD98hc^ from whole body to single-cell resolution. Our novel approach combining fluorescent tagging of molecules with FACS-sorting and single cell RNAseq reveals that ATV distributes robustly to numerous parenchymal cell types and blood-brain barrier endothelial cells, whereas standard antibody localization was limited to choroid plexus epithelial cells, meningeal, perivascular cells. At an organ level, we leveraged the enhanced 3D imaging capabilities of light sheet fluorescence microscopy and find distinct organ-specific biodistribution of ATV^TfR^, ATV^CD98hc^, and control IgG throughout the whole mouse body. Finally, 3D imaging in tissue-cleared cynomolgus monkey brains reveal widespread and enhanced distribution of ATV^TfR^ throughout the superficial and deep regions of both the brain and spinal cord. Altogether, these data shed further light on how the pathways governing IgG uptake and distribution enables broad delivery with ATV but severely limits distribution of standard antibodies in the CNS of both rodents and primates.

The peripheral distribution of each ATV platform was largely consistent with previous reports using more traditional methods ^1,2^, but the unbiased nature of the 3D whole-body imaging methodology reported here provide a more wholistic view of biodistribution and reveals important new insights for ATV^CD98hc^, including previously unappreciated localization to the lacrimal glands, parotid gland, and sciatic nerve. These findings from the present study highlights the potential for ATV^TfR^ and ATV^CD98hc^ to treat CNS diseases in which peripheral disease is also present, as well as provide additional biological considerations for platform-therapeutic pairings where there is a desire to target or avoid certain organs.

The contrast between TV-enabled and standard IgG biodistribution reveals several major limitations for standard antibody therapeutics for CNS diseases. Both our IHC and scRNA seq data strongly suggest standard non-BBB targeted IgG has a predominant route into the brain via the CSF, and not across the BBB. These data bear remarkable similarity to previously reported patterns for endogenous IgG ^13–15^, as well as intracisternal administered IgG ^16,57^. This plasma-to-CSF-to-brain trafficking route dominates for systemically administered IgG, with lowest concentrations in the deeper brain regions far from the ventricular and pial brain surfaces. Such patterns are expected to ultimately result in inefficient therapeutic target engagement, particularly in species with larger brain volumes ^63^. This distribution pattern may account for the typically higher exposures of standard antibodies detected in the CSF compared to brain parenchyma. Furthermore, this exposure profile (CSF IgG > CNS IgG) under normal conditions is also consistent with recognized differences in immune privilege between the CSF (a compartment associated with tissue graft rejection) and CNS parenchyma (a compartment associated with significantly blunted immune responses) that have been described for decades^64^.

The cellular distribution profile of control IgG is consistent with the interpretation that non-BBB targeted antibodies predominantly enter the CSF compartment from the circulation, rather than cross the BBB. Strikingly, less than 3% of BECs and pericytes had any control IgG signal, which may represent low levels of non-specific association with the cell surface facing the blood. Importantly, control IgG yielded almost no association with other parenchymal CNS cells. Rather, the cells with the highest control IgG signal were located at the inner and outer blood-CSF barriers (CPECs and fibroblasts type 1, including arachnoid barrier cells), suggesting these cells are likely involved in blood-to-CSF transport of standard IgG. This distribution agrees well with imaging results showing strongest signal in the choroid plexus and leptomeninges. Additionally, the perivascular signal on surface vessels and penetrating vessels points to subsequent CSF-to-brain entry of control IgG, supported by the cellular data showing localization with type 2 fibroblasts (including pial and perivascular fibroblasts) as well as perivascular lymphocytes and macrophages. This pattern of leptomeningeal and perivascular signal strikingly matches well with that previously reported for exogenous antibodies administered into the CSF as well as that of endogenous serum proteins, further supporting a mechanism of circulating IgG distribution first into the CSF and then into brain ^14,17^. Similar biodistribution patterns have also been observed when comparing intrathecal delivery of antisense oligonucleotides versus intravenous delivery of ATV^TfR^-enabled ASO ^33^, suggesting limited brain biodistribution likely poses a consistent challenge to other non-protein therapeutics when not enabled by a BBB-targeting platform. Importantly, even when bulk brain exposures were made equivalent by increasing the dose of control IgG by almost 7-fold over that of ATV^TfR^, IgG biodistribution to the parenchyma remained severely limited. Together, these observations underscore the critical importance of the brain entry route in shaping robust brain cellular biodistribution, with BBB entry greatly favored over CSF entry if the goal is to obtain widespread biodistribution to both superficial and deeper brain.

Consistent with active trafficking across the BBB as the predominant route of entry into the brain, ATV^TfR^ and ATV^CD98hc^ are highly localized to arteriovenous endothelial cells with lower levels in arterial compared to venous and capillary BECs. This pattern is also consistent with previously reported zonal expression of TfR and CD98hc (i.e., highest in capillaries and post-capillary venules) and direct blood-to-brain transport across the BECs ^53,62^. These distribution patterns translated to the cynomolgus monkey, where we observe greater ATV^TfR^ signal in smaller diameter capillaries and venules compared to larger diameter arteries. Each ATV also exhibit a unique cellular distribution profile to other CNS cells. ATV^TfR^ highly localizes to neurons, consistent with previous reports of brain TfR expression ^1^. Interestingly, ATV^TfR^ localizes to cell types that have not been previously appreciated, including astrocytes, and to a lower extent, to microglia and oligodendrocytes. ATV^CD98hc^ localizes to both astrocytes and microglia but no detectable localization was found with neurons (albeit in a limited sampling) or oligodendrocytes. Whole brain 3D imaging highlighted enhanced, widespread brain distribution for both ATV^TfR^ and ATV^CD98hc^ compared to control IgG. In particular, we noted several brain regions implicated in neurodegenerative diseases had relatively higher exposures (e.g., cortex, hippocampus, caudoputamen, thalamus) and may further benefit from ATV^TfR^ or ATV^CD98hc^ based drug delivery for these diseases. Broad distribution was also revealed in the spinal cords of mice, particularly in the more highly vascularized spinal grey matter, which was also observed in the cynomolgus monkey for ATV^TfR^.

Although this work provides important insights, the biodistribution of ATV-enabled therapeutic molecules will need to be further assessed independently, as both CNS cell-specific as well as peripheral distribution will be driven not only by TfR and CD98hc binding, but also by the therapeutic cargo (e.g. Fab targets, enzymes, ASOs, etc). Additionally, TV binding affinity, dose, and timepoint will likley all impact the distribution of molecules both in brain and periphery. While these results are relevant for the fundamental understanding of receptor-mediated BBB transport, each BBB technology will need to be evaluated separately, given that differences in architecture of the molecules could have significant impacts on CNS exposure and distribution. Additionally, the route of entry of standard IgG may have additional important therapeutic implications that warrants further investigation. For example, it will be interesting for future studies to evaluate whether ATV-enabled delivery of anti-amyloid antibodies across the BBB can reduce engagement with the vascular amyloid commonly present on leptomeningeal arteries and perivascular drainage pathways ^54,55,65^. Avoiding these pathways with a BBB-dominant route of entry into the CNS could potentially mitigate risks such as immunotherapy-induced vascular inflammation and microhemorrhages^56,66,67^. Nevertheless, our results provide a thorough foundational understanding of the brain cell types and tissues targeted by each TV platform, which will enable optimal platform selection to efficiently drive desired distribution and target engagement profiles of a variety of TV-enabled therapeutics for neurological diseases.

## Acknowledgements

The authors would like to thank Rene Meisner for her input on the anatomical interpretation of whole-body tissue clearing imaging results; Shan V. Andrews, Alexander Schuth, Mitesh Shridhar, and Ryan Watts for providing helpful comments on the manuscript, Butch Benitez, Kevin Rebadulla, and Dominic Sobrepena for their help with mice care at Denali Therapeutics.

## Author contributions

**Conceptualization** N. K., M. E. P., R.T., M.E.K.C., Y.J.Y.Z. **Data generation and or analysis** N.K., M.E.P., C.B.D., D.J., D.T., M.I.T., M.N., C.H., G.L.D.M., L.S., M.J.S., D.C., R.C., K.C., J.C., A. C., Y. R.C., J.C.D., J.D., D.K., H.K., A.L., E.L., A.M., E.R., T.S., M.T., K.X., Y.Z., M.E.K.C. **Writing and main edits** N.K., M.E.P., R.G.T., M.E.K.C., Y.J.Y.Z. **Supervision** J.L., A.E., R.G.T., M.E.K.C., Y.J.Y.Z.

## Methods

### Animal care

All animal procedures were performed in adherence to ethical regulations and protocols approved by Denali Therapeutic Institutional Animal Care and Use Committee. Mice were housed under a 12h light/dark cycle and had access to water and a standard rodent diet (LabDiet 5LG4, Irradiated) *ad libitum*. Temperature and humidity in all animal rooms were monitored daily by Thermo Scientific^TM^ InSight. The normal temperature range was 18.3-23.3°C and the normal humidity range was 30–70%.

### *In vivo* mouse studies

Mice used in this experiment included C57BL/6J, TfR ^mu/hu^ KI, and CD98hc^mu/hu^ KI (on a C57Bl6J background) (The Jackson Laboratory). For the whole body biodistribution and brain imaging experiments, 12-week old male mice were used. For the FACS-scRNA-seq experiment, 12-week old mixed sex mice were used with 2 males and 2 females per group. For the brain concentration matching experiments, female TfR^mu/hu^ KI mice were used at 18-19 weeks of age. Mice were IV dosed via the tail vein with the test articles and then anesthetized deeply via IP injection of 2.5% Avertin at the terminal timepoints. For PK studies, mice were transcardially perfused with ice cold PBS at a rate of 5 mL/min and brains were collected.

### TV engineering

ATV^TfR^ and ATV^CD98hc^ platforms were engineered as previously described using a huIgG1 backbone ^1,2^. For the mouse studies, we used a monovalent form of ATV^TfR^ clone 35.23.2 or clone 35.23.4^1^ and the monovalent form of ATV^CD98h^ clone LLB6.8 ^2^. For the cynomolgus monkey studies, we used a monovalent form of ATV^TfR^ clone that binds cynomolgus monkey TfR^33^. To be able to dissect the contribution of platform alone to biodistribution, our control huIgG1 antibody, ATV^TfR^, and ATV^CD98hc^ were engineered with bivalent non-targeting control Fabs that bind a small molecule, the hapten dinitrophenol (DNP), not present *in vivo*^68^. Additionally, we introduced the L234A/L235A/P329G (LALAPG) mutations to the Fc region which abolish FcγR and C1q binding and consequently eliminate effector function activity ^44^.

### Fluorescent antibody conjugation

To fluorescently conjugate our antibodies, we used the Thermo Fisher Scientific SiteClick Labeling Kits (S10911, S10901) which allow the attachment of Alexa-647 to the heavy chain N-linked glycans. The main advantage of this approach is that the conjugation is far from the antigen-binding domain and the TV location which reduces the potential for interference with molecule biochemical properties. On average we achieved a degree of labelling between 1-2 AF647 fluorophores per antibody. This was determined by the absorbance of the fluorophore and antibody on the nanodrop and followed by the calculation using the kit’s recommendation. Quality control steps including *in vitro* (cell binding and SPR (Surface Plasmon Resonance)) and *in vivo* (wild-type plasma clearance and brain uptake) were conducted to ensure the fluorophore does not interfere with the binding and biodistribution properties of the ATV platform *(data not shown)*.

### Brain huIgG quantification

The left side of the frontal brains were fresh-frozen and homogenized using a Qiagen TissueLyser with 5 mm steal beads for 6 min at 30 Hz in 10x volume / tissue weight of lysis buffer containing 1% NP-40 in PBS with protease inhibitors. Homogenate was centrifuged at 14,000 rpm for 20 min at 4°C and supernatant collected. Brain lysates were diluted 1:2 and 1:20 for analysis of huIgG concentration.

HuIgG concentrations were quantified using a generic anti-human IgG sandwich-format ELISA. Briefly, plates were coated overnight at 4 °C with donkey anti-human IgG (JIR #709-006-098) at 1μg/mL in sodium bicarbonate solution (Sigma #C3041-50CAP) with gentle agitation. Plates were then washed 3x with wash buffer (PBS + 0.05% Tween 20). Assay standards and samples were diluted in PBS + 0.05% Tween 20 and 1% BSA. Standard curve preparation ranged from 0.41 to 1500 ng/mL or 0.003 to 10 nM (BLQ < 0.03 nM). Standards and diluted samples were incubated with agitation for 2 h at room temperature. After incubation, plates were washed 3× with wash buffer. The detection antibody, goat anti-human IgG (JIR #109-036-098), was diluted in blocking buffer (PBS + 0.05% Tween-20 + 5% BSA) to a final concentration of 0.02 μg/mL and plates were incubated with agitation for 1 h at room temperature. After a final 3× wash, plates were developed by adding TMB substrate and incubated for 5–10 min. Reaction was quenched by adding 4 N H_2_SO_4_ and read using 450 nm absorbance.

### Mouse immunohistochemistry

For the experiment involving dose-matched 2D imaging, after perfusing the mice with PBS as described above, the hemibrains were drop fixed in 4% PFA overnight. Sagittal brain sections (40 μm) were cut using a microtome (MultiBrain® Technology by NeuroScience Associates), blocked in 5% BSA + 0.3% Triton X-100. Sections were mounted in Prolong glass (Thermofisher P36984) before imaging at 20x magnification using a widefield epifluorescence slide scanner (Axio Scan Z1; Carl Zeiss; 20x/0.8 NA air objective; each channel was acquired independently with the appropriate filter cube for that fluorophore).

For the experiment involving brain-concentration-matched 2D imaging, perfused hemibrains were immersion fixed in 4% PFA for approximately 24 h at 4C before transferring to PBS + 0.01% sodium azide followed by sectioning sagittal at 40 μm thickness (MultiBrain® Technology by NeuroScience Associates). Serial sections were stained free-floating by incubating in blocking buffer (PBS/TBS + 1% BSA + 1x fish gelatin (BioWorld 21761058) + 0.5% triton X-100 + 0.1% sodium azide) at room temperature for 2 h, incubation with primary antibodies (in PBS + 1% BSA + 0.3% triton x-100 + 0.01% sodium azide) overnight at 4C, three fifteen minute washes in PBS + 0.3% triton X-100, incubation in secondary antibody (in PBS + 1% BSA + 0.3% triton x-100 + 0.01% sodium azide with DAPI) for 4-5 h, and three 15 min washes in PBS + 0.3% triton x-100. Sections were mounted onto slides and sealed with a coverslip with Prolong Glass (Invitrogen, P36984) before imaging at 20x magnification using a widefield epifluorescence slide scanner (Axio Scan Z1; Carl Zeiss; 20x/0.8 NA air objective; each channel was acquired independently with the appropriate filter cube for that fluorophore).

### Mouse 3D brain tissue clearing and imaging

After 24 h of dosing mice with 100 mg/kg control IgG or 15 mg/kg ATV^TfR^ through tail vein IV injection (brain-concentration-matched study), mice were deeply anesthetized via IP injection of 2.5% Avertin and immediately transcardially perfused using ice cold PBS containing 5 mg/L LEL tomato lectin (DyLight 649 conjugated; Invitrogen, L32472) at a flow rate of 2.5 mL/min for 6 minutes, followed by perfusion of room temperature 4% PFA for 25 minutes. Spinal cords and brains were extracted, the brains bisected, and each tissue post-fixed in 4% PFA for 24 hours, rinsed in PBS, and then transferred to PBS + 0.01% sodium azide and stored at 4C, protected from light throughout the process. One animal in the control IgG group was not well fixed (body not rigid) due to an air bubble in the line during perfusion-fixation and was excluded from further analysis.

Samples were brought to RT and washed in 1xPBS before being dehydrated in a 200-proof methanol/milliq water gradient of 20%, 40%, 60%, 80% and 100% at 1 hour while on a rocker at gentle speed. After washing once more in 100% methanol at room temperature, samples were incubated overnight in 67% dichloromethane/33% methanol, after which they were washed twice in 100% methanol for 30 minutes (RT), cooled to 4C and bleached in chilled fresh 5% hydrogen peroxide in methanol overnight at 4C. After rehydration in a room-temperature methanol/PBST series (1 hour each at 80%, 60%, 40%, 20% methanol in PBS with 0.2% TX-100), samples were washed in PBST twice for an hour before incubating in permeabilization solution (2.3% glycine, 20% DMSO, 0.02% sodium azide in PBST) at 37C for 4 days. Once permeabilized, samples were blocked at 37C for 3 days in a solution of 6% donkey serum, 10% DMSO, 0.02% sodium azide, and 0.2% triton x-100 in PBS. After blocking, samples were incubated with primary antibody in a buffer of 0.2% gelatin, 0.5% trionx100 and 0.1% saponin in PBS for 14 days at room temperature, while being rocked gently. After washing (in PBS with 0.1% saponin, 0.2% tween20, and 0.1% of a 10mg/ml heparin solution) for 1x10 min, 1x20 min, 1x30 min, 1x1 hr, 1x1.5hr (at RT) and 1x 3 days (at 4C), samples were incubated in secondary antibody for 7 days. The secondary incubation took place in antibody dilution buffer (0.2% tween20, 0.1% of 10mg/ml heparin stock solution, 0.1% saponin, 0.5% tritonX100, 0.2% gelatin) at RT with gentle rocking. Samples were then washed again (as after primary antibody incubation), and dehydrated in a methanol/milliq water gradient as before. After a final overnight incubation in 100% methanol, samples were washed for an hour in fresh 100% methanol and incubated in 66%DCM (Dichloromethane)/33% methanol at room temperature and finally washed in 100% DCM 15 minutes twice (with shaking) to remove traces of methanol. Stained and delipidated tissue samples were cleared in DiBenzyl Ether in airtight vials for 24 hours, and in ECi for another 24 hours (RT and protected from light) before imaging in ECi. Antibodies used were donkey anti-huIgG/DL800 (1:500 Rockland 605-745-125), Goat anti-CD31 (1:300 RD systems AF3628), Goat anti-Podocalyxin (1:1000 RD systems AF1556) and donkey anti-goat Cy3 (1:500 Jackson ImmunoResearch 705-166-147.)

Fully cleared samples were imaged in the Miltenyi Ultramicroscope blaze with the 1x objective (1x zoom), and again with 12x objective (1x zoom). For both imaging runs, photographs were taken with 0.186 numerical aperture, with two-sided illumination, adaptive focus and adaptive blending. All images of one magnification were taken with the same optimized settings. 1x images were taken at 70% sheetwidth, at 3.86µM steps. Excitation (Emission) as follows: 785(805): 10%, 640 (680) 7%, 561(620) 5%, and 488 (525) 40%. 12x images (of the medial branch of the circle of Willis) were taken at 50% sheetwidth, at 0.3µM steps. Excitation (Emission) as follows: 785(805): 8%, 640 (680) 9%, 561(620) 6%, and 488 (525) 77%.

### FACS-scRNA-seq

For each treatment group (Ctrl IgG, ATV^TfR^, ATV^CD98hc^), 2 male and 2 female mice were perfused with 1x PBS and their whole brain were collected (excluding olfactory bulbs as well as cerebellum, pons, and medulla). For downstream processing, brains from 1 male and 1 female per treatment were pooled and processed together for single cell dissociation in order to reduce the number and cost of single cell RNA-seq libraries. Cells from individual animals were then demultiplexed based on sex-specific gene expression allowing us to analyze cells from all 4 animals per treatment group.

To dissociate the brains into single cells we used the Neural Tissue Dissociation Kit (Miltenyi Biotec, cat. #130-092-628) as previously described ^69^. Briefly, the brains were chopped using a razor, resuspended with HBSS, and centrifuged. The pellet was resuspended and incubated with the enzyme P mix at 37°C for 10min along with the transcriptional and translational inhibitors Actinomycin D (Cell Signaling Technology, cat. #15021, 5uM final concentration) and Anisomycin (Cell Signaling Technology, cat. #2222, 2uM final concentration) respectively. Samples were then homogenized by trituration using a 5ml serological pipette and incubated with Enzyme A at 37°C for another 10min. Following incubation, the brains were triturated with a 2ml serological pipette, washed with DPBS, and passed through a 100uM filters and spun at 300xg 10min. To remove the excess myelin, cells were resuspended in 0.9M sucrose and spun at 850g for 35min. Cells were then washed with FACS buffer (1x PBS, 1% BSA, 2mM EDTA) and stained with SYTOX™ blue (ThermoFisher) to distinguish live cells.

On the FACS sorter (BD FACSAria™ III Cell Sorter), the gates were set to capture single live cells, and using the naïve mouse sample the Alexa-647 positive gate was drawn. We used ATV^TfR^ samples to draw the gates for the low, mid, and high samples (ensuring no overlap) such that each gate would capture 33% of the cells from the AF647-positive gate for that group. To have a fair comparison across treatments, these same gates were applied to all other groups. Additionally, since the AF647-negative bin accounts for 99% of the cells in the IgG control group, sorting cells from this bin for the same duration of time as the low, mid, and high AF647-bins (total of ∼1% for the IgG control group) would have exceeded the capacity of the collection tubes for the AF647-ve bin. Thus, we decided to sort only 60,000 cells for the AF647-ve bin for all treatment groups and record the time required to obtain this cell number. Afterward, we simultaneously sorted the cells from the AF647-low, -mid, and -high bins as well as recorded the time required. Then to obtain the theoretical number for the AF647-ve bin that we would have obtained, we extrapolated this number from the total time it took to sort the AF647-low, mid, and high bins for that group (further described in **Single-cell RNA-seq data processing and analysis**).

### Single-cell RNA-seq library construction and sequencing

Six single-cell suspensions of pooled male/female dissociated brain tissues were FACS sorted into four different bins based on AF647 signal intensity resulting in 24 samples for single-cell capture. Up to 60,000 cells were sorted into each 1.5 mL centrifuge tube containing the 24 samples consisting of the 3 treatment groups with 4 bins each and 2 pooled animals per treatment group (1 male, 1 female). To concentrate the single-cell suspensions, each sample was pelleted by centrifugation at 200xg for 10 minutes in a swing bucket rotor at 4°C, and supernatant was removed leaving only 43.3 µL volume for resuspension. The entire volume of cells was then added to the RT master mix and loaded onto Chromium Next GEM Chip G microfluidic channels per manufacturer’s protocol (10X Genomics, CG000315 Rev C). One sample (IgG1-bin 0) was lost due to a fluidic wetting issue, leaving 23 samples for library preparation and sequencing analysis.

Post GEM–RT Cleanup was performed using a Dynabeads cleanup mix via magnetic purification, and cDNA was generated using 12 cycles of amplification followed by cDNA purification using SPRIselect reagent. A quarter volume of the cDNA generated was used as input for 3’ gene expression library construction starting with fragmentation, end repair and A-tailing, followed by Illumina adaptor ligation and sample index PCR, with SPRIselect bead purification performed in between each step.

Library quantity and quality were assessed with High Sensitivity D1000 ScreenTapes (Agilent 5067-5584) and then pooled in equimolar ratios for shallow sequencing on an Illumina MiSeq Reagent Kit v3 (Illumina, MS-102-3001) to determine cell capture rate. With the cell capture number information from the MiSeq sequencing result, libraries were pooled targeting 20,000 reads per captured cell. RNA-seq data for analysis was generated on an Illumina NovaSeq 6000 instrument, S2 cartridge, Paired end (28X10X10X90) by SeqMatic (Fremont, CA, USA).

### Single-cell RNA-seq data processing and analysis

After sequencing, fastq files were processed using Cell Ranger (v7.1.0). To increase the number of samples without generating additional libraries, each library consisted of one male and one female mouse which could then be computationally demultiplexed using gene expression data. For every cell, male/female identity was predicted using a random forest classifier trained on mouse single-cell data from the Linnarsson Adolescent Mouse Brain Atlas ^70^. The model was trained on 6 genes that have strong differential expression between males and females: *Xist*, *Tsix*, *Ddx3y*, *Kdm5d*, *Uty*, and *Eif2s3y*. After applying a 90% confidence threshold, the model predicted sex with ∼99% accuracy. This accuracy was supported by observed separation of X and Y chromosome gene expression between predicted male and female mice (**Supplementary** Fig 2a). Normalization, variable feature selection, PCA, UMAP dimensionality reduction, and louvain clustering were all carried out in R using Seurat v4 ^71^. Broad cell types were predicted using the Seurat label transfer methodology and an internally-generated single-nucleus atlas of mouse brain cell types. Predictive cell typing was supported by observation of top marker genes in each predicted class of cells (**Supplementary** Fig 2b). Following cell typing, putative doublets were identified and removed using the scDblFinder package in R ^72^. Fine cell typing of immune and endothelial populations was performed using iterative subclustering and manual cell typing through marker gene analysis. Briefly, the populations were isolated and clustered, obvious doublet clusters were removed, and cluster identities were mapped to known cell types by examination of canonical marker gene expression. These detailed cell type labels were projected back onto the full object as seen in Figure 2.

Because FACS sorting had to be run for much longer to capture the A647+ gates, the raw capture rates between the A647-gate and the A647+ gates could not be compared directly. Therefore, cell type distributions across these gates were calculated using a correction factor applied to the A647- samples. For each mouse, a correction factor was calculated by dividing the sort time for A647+ gates by the sort time for A647-. The raw number of cells in the A647-sample was then multiplied by this correction factor, representing a hypothetical A647-capture rate that would have been achieved if the FACS had been run for as long as it was for the A647+ gates. In this way, interpretable percentages of labeled and unlabeled cells could be calculated for each cell type. All reported percentages through figure 2 and figure 3 use these adjusted values.

Pseudobulk expression values were computed for each sample by summing single-cell expression within cell types across all samples using the AggregateAcrossCells function in the scuttle R package. Log CPMs (counts per million) were calculated using a pseudocount of 1.

### Whole body tissue clearing

Mice were transcardially perfused with ice cold PBS at a rate of 4 mL/min followed by 20 min perfusion with 4% PFA. They were then post-fixed for 12hrs with 4% PFA then washed with PBS and stored in PBS with 0.05% sodium azide until ready for shipment. Upon arrival to Deep Piction, they were skinned, washed in PBS and set up for whole-body perfusion as described in the vDISCO protocol, omitting the nanobody staining step ^42^. In brief, the mice were stored in a glass staining jar and a stainless-steel oral gavage needle was attached to the left ventricle of the heart and secured with superglue. Subsequently, the mice were perfused with the following solutions at room temperature: PBS (1x 12h), CUBIC-I solution for decolorization (4x 12h), PBS (3x 3h), 20% EDTA for decalcification (3d), PBS (3x 3h), Permeabilization solution (3.5d). Subsequently, the perfusion needle was removed, and the stomach and intestines were cleaned of chyme and feces by small incisions every 2-3 cm and careful rinsing with PBS via a syringe. The mice were then washed 2x 2h in PBS and dehydrated at RT on a shaker in a series of Tetrahydrofuran (THF) /dH2O mixes with increasing THF concentrations as follows: 50% THF (12h), 70% THF (12h), 90% THF (12h), 100% THF (2x 12h). They were then dilapidated in Dichloromethane (DCM) for 6h and refractive index matched in a 1:2 mixture of Benzyl Alcohol / Benzyl Benzoate (BABB). We replaced the BABB after the first 12h for better clearing performance.

### Whole body tissue clearing imaging and processing

The mice were scanned in a customized Ultramicroscope Blaze (Miltenyi Biotec, Germany) outfitted with a custom whole-mouse imaging chamber and a custom stage. Samples were imaged with a 1.1x objective optimized for BABB imaging (LaVision-Miltenyi BioTec MI PLAN ×1.1/0.1 NA, WD 17 mm) and recorded with a scientific sCMOS camera (Miltenyi Biotec, resolution 2048x2048 px / frame, 16-bit depth, effective pixel size 5.8µm/px). Samples were superglued to the sample holder and first imaged along the ventral side with z-step size of 6 µm/step (ca 1700 z-planes) and a 3x9 tiling layout in X/Y (25% overlap), The light-sheet thickness was set to 11.4µm (sheet NA: 0.03) and sheet width of 100%, which evenly covered the entire field of view of the 1.1x objective. The exposure time was set to 80ms/frame for imaging of the background tissue autofluorescence using a 488nm laser (emission filters 525/50, 12% laser power) and AlexaFluor-647 conjugated antibody signal with a 640nm laser (emission filters 680/30, 14% laser power). Following ventral imaging, mice were flipped to the dorsal position, re-glued, and the dorsal half of the animal was imaged using the same parameters.

Imaged tiles were stitched with Fiji (ImageJ) Stitch Sequence of Grids of Images function. For the organ-wise region of interest (ROI) analysis, the whole-body image stack (signal channel only) was loaded as a virtual stack in Fij and placed a set of markers on the respective organs (minimum two per organ). Regions Of Interest (ROIs) were then processed by cutting out a 10x10x10 voxel cube centered around the marker. The median, standard deviation, minimum and maximum values were first calculated from all voxels. These were then grouped by ROI type and treatment group, normalized the means to the mean of the control condition, and log-transformed the results.

In addition, the whole-body image stacks were segmented into organs of interest, brain, kidney and lungs, using a custom machine learning pipeline.

### Mouse brain regional analysis

Segmented brains were resampled from their initial resolution of 5.80 μm x 5.80 μm x 6.00 μm to 10 μm isotropic volumes with 3^rd^ order b-spline interpolation using the ResampleImageFilter in the Insight Toolkit (ITK) version 5.3.0 ^73^. Using this resampled volume, an image pyramid with isotropic resolutions 10 μm, 25 μm, 50 μm, and 100 μm was built for each brain image, corresponding to the voxel pitches of the Allen Institute of Brain Science common coordinate framework version 3 brain atlas (CCFv3) ^74^.

The CCFv3 atlas defines a nested hierarchy of over 1,300 brain regions, with the most finely delineated layers at or below the resolution of the segmented whole brain data. To enhance the robustness of our region-based quantification, we collapsed the CCFv3 annotations into 14 high-level brain regions by combining the parent region mask(s) and all child region mask(s) into a single region of interest (ROI) as described in **Supplementary Table 1**.

Individual brain volumes were registered in stages to the Nissl-stained template using the ElastixImageFilter class in itk-elastix v0.17.1 ^75^. iDISCO cleared brains were uniformly flatter along the dorsal-ventral (D/V) axis compared to the template image, so the 25 μm brain volumes were each expanded by 50% along the D/V axis, then rigidly registered to the 25 μm Nissl-stained atlas image using parameters given by the GetDefaultParameterMap(“rigid”) function in itk-elastix, coarsely aligning the two in space. Next, the rigidly aligned brains were further registered to the Nissl template with an affine transform using parameters from the GetDefaultParameterMap(“affine”) function. Finally, each brainwas nonlinearly warped with a B-spline warp using parameters from the GetDefaultParameterMap(“bspline”) function, except with a final grid spacing of 1,600 μm, 9 rounds of optimization, and a grid spacing schedule of 8x, 8x, 8x, 4x, 4x, 4x, 2x, 2x, 2x for each round respectively. Warped brains were visually overlaid on the atlas Nissl template image to assess registration quality using napari v0.4.19) ^76^. While gross region alignment was good across all brains, fine structures were visibly misaligned. To improve alignment of fine structures, all 9 b-spline warped brain images and the 25 μm Nissl atlas template were simultaneously aligned using the parameters from the GetDefaultParameterMap(“groupwise”) function, except with a final grid spacing of 800 μm, an increased maximum number of iterations to 512 steps, and a 10-step grid spacing schedule with 2 steps each of 32x, 24x, 16x 12x and 8x optimization. To transfer ROIs onto the brains, both the high level merged ROIs described above, and the lowest level annotations for each voxel of the 25 μm Allen Brain Atlas were warped through the rigid, affine, b-spline and groupwise sequence using the TransformixImageFilter with a final b-spline interpolation order of 0 (nearest neighbor interpolation)

A brain mask was defined on the 25 μm isotropic fluorescence image by thresholding the image to include the entire visible brain volume as well as any surface vessels. Voxel-wise brain depth was then quantified inside this brain mask using the distance_transform_edt function from scipy v1.11.4 ^77^. Mean intensity values were quantified as the mean huIgG intensity within the intersection of each warped brain atlas ROI and the whole brain mask. For hierarchically defined brain regions, mean intensity was calculated for all voxels annotated with that region id or any descendent region ids. Normalized mean intensity was calculated for each region by dividing the regional mean intensity by the regional mean intensity for Brain Atlas region id #8, basic cell groups and regions. Annotations from the fused atlas were warped back onto the original image space using the numerical inverses of the bspline, affine, then rigid transforms above, then overlaid onto the 10 μm isotropic brains in native space. Depth-wise mean intensity was calculated by binning depth from surface into 10 μm bins, then calculating mean signal within the intersection of each inverse warped brain atlas ROI and the depth bin.

### *In vivo* cynomolgus monkey study

Female cynomolgus monkeys (2.6-3.6 years old, Macaca fascicularis, n = 1/group; Charles River Laboratories) were housed at 64–84 °F, 30–70% humidity. Housing set-up is as specified in the USDA Animal Welfare Act (Code of Federal Regulations, Title 9) and as described in the Guide for the Care and Use of Laboratory Animals. Cynomolgus monkeys were IV administered with a single 23 mg/kg dose of appropriate test article, 2 days post-dose the monkeys were anesthetized with ketamine HCl combined with isoflurane/O2 and perfused with 4ml phosphate buffer saline (PBS) containing heparin, followed by 12.5 mg/kg of WGA-Alexa555 (wheat germ agglutinin) 1.6 mg/ml to label blood vessels, and then 3ml of 1x PBS. After euthanasia, the brain and spinal cord were collected and drop fixed in 4% PFA overnight and then stored in 1xPBS with sodium azide.

### Cynomolgus monkey brain and spinal cord clearing

Hemi-brains were washed twice in a 10x volume of 1xPBS for a total of 24 hours at room temperature with gentle rocking, protected from light. The tissue was then dehydrated through a series of 200 proof fresh ethanol (LC/MS grade) and milliq water solutions: first at 50% ethanol for 48 hours (37C), followed by 4 days at 70% ethanol (RT), 48 hours at 80% (37C), 48 hours at 90% (37C), 72 hours at 100% (RT), and finally 48 hours at fresh 100% (37C). After dehydration, tissue was incubated in a mixture of 50% EtOH and 50% THF for 24 hours at 37C, followed by a 4-day incubation in 100% THF (37C), and finally a 6-day incubation in 100% DCM (37C).

Hemi-brains were then cleared in 99.9% ECi at RT in the dark without agitation for 3 days, before being switched to fresh ECi and allowed to clear further for another 5 days before imaging in ECi in the Miltenyi Blaze Ultramicroscope.

Spinal cord segments were chosen to be comparable between the two animals, taken from the first three, non-enlargement cervical spinal segments. After bringing samples to room temperature, tissue was washed once in PBS for an hour on gentle rocking at RT before dehydrating in similar manner to the brain samples, in 15ml of an ethanol/water series. First, tissue was incubated in 50% ethanol for 3 hours (RT), followed by 70% ethanol overnight (RT), 80% for 3 hours (RT), 90% 3 hours (RT), 100% overnight (37C), and finally fresh 100% for 3 hours (37C). After dehydration, tissue was moved to a 10ml glass vial filled completely with 50% EtOH & 50% THF, and rocked gently for 3 hours at 37C, before incubating in 100% THF overnight (37C). The following day, samples were rocked gently in 100% THF for 3 hours (37C) before moving to 100% Dichloromethane overnight (37C). Finally, delipidated cord segments were cleared in 99% Ethyl cinnamate (ECi) for 3 days, followed by switching to fresh ECi for 24 hours before imaging in ECi in the Miltenyi Blaze Ultramicroscope.

Samples from the two animals were treated identically and imaged using the same parameters, the details of which varied with the region of interest (hemi-brain or specific ROI), tissue (hemi-brain or spinal cord), and objective used.

### Cynomolgus monkey brain imaging

All samples were imaged in ethyl cinammate (ECi), with a retail Ultramicroscope Blaze microscope using the standard 2048x2048 px / frame 16-bit depth sCMOS camera, with the exposure time set to 650ms.For overview scans, the samples were imaged using the standard organic-solvent-optimized 1.1x objective and a 0.6x zoom-out setting in the ImSpector (v7.5.3) software using the 640 laser (emission filter 680/30 16% laser power) in a 2x4 tiling grid with 35% overlap and dual-sided illumination. The samples were imaged sagitally, first glued on the medial plane so that the lateral cortex was closest to the objective (lateral scan), then they were repositioned so that the medial sulcus was closest to the objective (medial scan). The z-step size was set to 13µm. Detailed volume imaging was done with the standard organic-solvent-optimized 4x objective without additional zoom in the software, by using the 561nm and 640nm lasers (emission filters 620/60 and 680/30 respectively and 14% laser power) in a 2x3 tiling grid with 33% overlap and single sided illumination. The z-stepping was set to 2.71µm. The samples were positioned in an oblique plane, glued on the medial plane so that the ventral hippocampal region was closest to the objective and the dorsal cortex was next to the sample holder bar, below the laser’s height.

### Cynomolgus monkey spinal cord imaging

All samples were imaged in ethyl cinnamate (ECi), with a retail Ultramicroscope Blaze microscope using the standard organic-solvent-optimized 4x objective and the 0.6x zoom-out setting in the ImSpector acquisition software. We used a standard 2048x2048 px / frame 16-bit depth sCMOS camera with 264ms and 650ms exposure times for the 561nm and 640nm lasers resapectively. We used the 561nm and 640nm lasers (emission filters 620/60 and 680/30 respectively and 14% laser power) for dual-sided illumination. The z-stepping was set to 2.71µm.

### Cynomolgus monkey vessel processing

The produced image stacks were stitched using TeraStitcher. The lateral and medial overview scans were fused together using Vision4D’s landmark-based Volume Fusion function to a single volume. The detailed scans were segmented using VesSAP’s vascular segmentation in the channel highlighting the brain blood vessels. This mask was then expanded using Fiji’s Maximum 3D filter using 2, 4 and 6 voxels in XYZ. To analyze the distribution of parenchymal intensity, we first removed the vascular mask from the 2vx expanded mask, resulting in a concentric shell surrounding the blood vessels. Then, we removed the 2vx mask from the 4vx mask, forming a larger shell, and then removed the 4vx shell from the 6vx mask to create the outermost shell surrounding the vasculature. These masks were applied to the signal channel only. Next, a 5-step tissue depth binning was produced in Fiji using polygon selections, each bin encompassing a 135µm (50vx) depth. The histogram of each vascular and parenchymal shell (in each depth bin) was extracted using the standard Fiji function and the weighted mean was compared.

To extract the local vessel radius, the vessel mask was first processed with an iterative quasi-3D hole closing filter. For each possible Z plane, the XY vessel mask was extracted then filtered using the remove small holes function from scikit-image with a maximum area threshold of 7,344µm^2^ (1000 px^2^). This filter was repeated for each possible Y plane filtering the XZ mask, followed by each possible X plane filtering the YZ mask. The vessel mask was next processed with a binary opening operation to remove axis-aligned stripe artifacts. The morphological skeleton was then calculated using a 3D skeletonization algorithm ^78^ and the distance from the edge of the hole filled vessel mask to the skeleton was calculated using a Euclidean distance transform ^79^ . Each point along the skeleton was assigned an index and the distance from the edge at that index was used as the estimate for the local radius at that point.

Each voxel of the hole-filled vessel mask was assigned to the nearest point on the vessel skeleton using watershed segmentation. This segmentation was then masked so only voxels in the original segmentation were considered, and any voxels not assigned to a corresponding point on the vessel skeleton were filtered (∼1.0-1.5% of the vessel mask). The mean huIgG was then calculated for each label in the segmentation, resulting in a measurement of radius and mean intensity at each point in the vessel skeleton. The radii were binned into 20 buckets from 2.71 µm (1 voxel) to 54.2 µm (20 voxels) and the mean intensity within each bin was calculated.

### Cynomolgus monkey brain atlas registration

1x magnification tiled scans of the cynomolgus brains were first stitched using TeraStitcher then converted to Imaris image pyramid volumes using ImarisFileConverter. The 2x down sampled volume was then loaded and flipped along the y-axis to bring it into rough alignment with the T1 weighted template image for the 24-month-old cynomolgus atlas ^80^. The brain volume was N4 corrected using a shrink factor of 32x, with 5 levels of correction and 50 iterations at each level. The corrected brain was thresholded followed by morphological filtering to remove mask fragments smaller than 0.00292 mm^3^ (500 vx) and to remove holes smaller than 0.292 mm^3^ (50,000 vx), then all values outside the brain mask were set to 0.

The T1 weighted template image was split along the midline to produce a hemibrain image, then trimmed 1.9 mm (5 voxels) on each side to match the approximate imaging area of the light sheet image. The template image was then upsampled, and the corrected brain image down sampled each to 0.25 mm x 0.25 mm x 0.25 mm isotropic using linear resampling with the ResampleImageFiler from ITK. The corrected brain image was then registered to the trimmed T1 template image using ANTsPy (antspyx v0.4.2)^81^. To calculate a robust initial transform, the corrected brain was first coarsely aligned to the template using the ‘Rigid’ transform preset, then this initial transform was further refined using the ‘Affine’ transform preset. The resulting affine initial transform was used to initialize the ‘SyNAggro’ transform preset with no additional refining affine iterations, SyN sampling of 128 per iteration, and a series of six levels of registration with 80, 80, 80, 40, 20, and 10 iterations per level respectively. This resulted in a set of forward and inverse affine and warp parameters mapping between the input volume and the atlas template space.

The SARM and CHARM annotations for level 2 of the cynomolgus 48-month atlas were transformed back to the 0.25 mm isotropic image space using the apply_transforms method from ANTsPy with the ‘genericLabel’ interpolator. The labeling was manually inspected on the original images using napari to ensure alignment with the light sheet anatomy. To enable comparison with the mouse atlas, the CHARM cortical parcellation was combined into a single “cortex” region, while the SARM labels were not modified. Mean intensity was calculated within each region inside the brain mask using the (UNCORRECTED|N4 CORRECTED) images.

### Statistical analysis

One-way ANOVAs were performed using Prism 9 software (GraphPad 9.5.1).

### Data Availability

The raw and processed scRNA-seq data are available via the Gene Expression Omnibus (GEO) under the accession number GSE262436.

### Code Availability

Manuscript code will be made available in Zenodo upon publication.

## Figure Legends

**Video 1:** Mouse whole-body IgG, ATV^TfR^ and ATV^CD98hc^ biodistribution

**Video 2:** Control IgG biodistribution in AI-segmented mouse brains

**Video 3:** ATV^TfR^ biodistribution in AI-segmented mouse brains

**Video 4:** ATV^CD98hc^ biodistribution in AI-segmented mouse brains

**Video 5:** Control IgG biodistribution cynomolgus monkey brains

**Video 6:** ATV^TfR^ biodistribution cynomolgus monkey brains

