## Supplemental Info for "Fc-engineered large molecules targeting blood-brain barrier transferrin receptor and CD98hc have distinct central nervous system and peripheral biodistribution compared to standard antibodies"

### Supplementary Figure 1

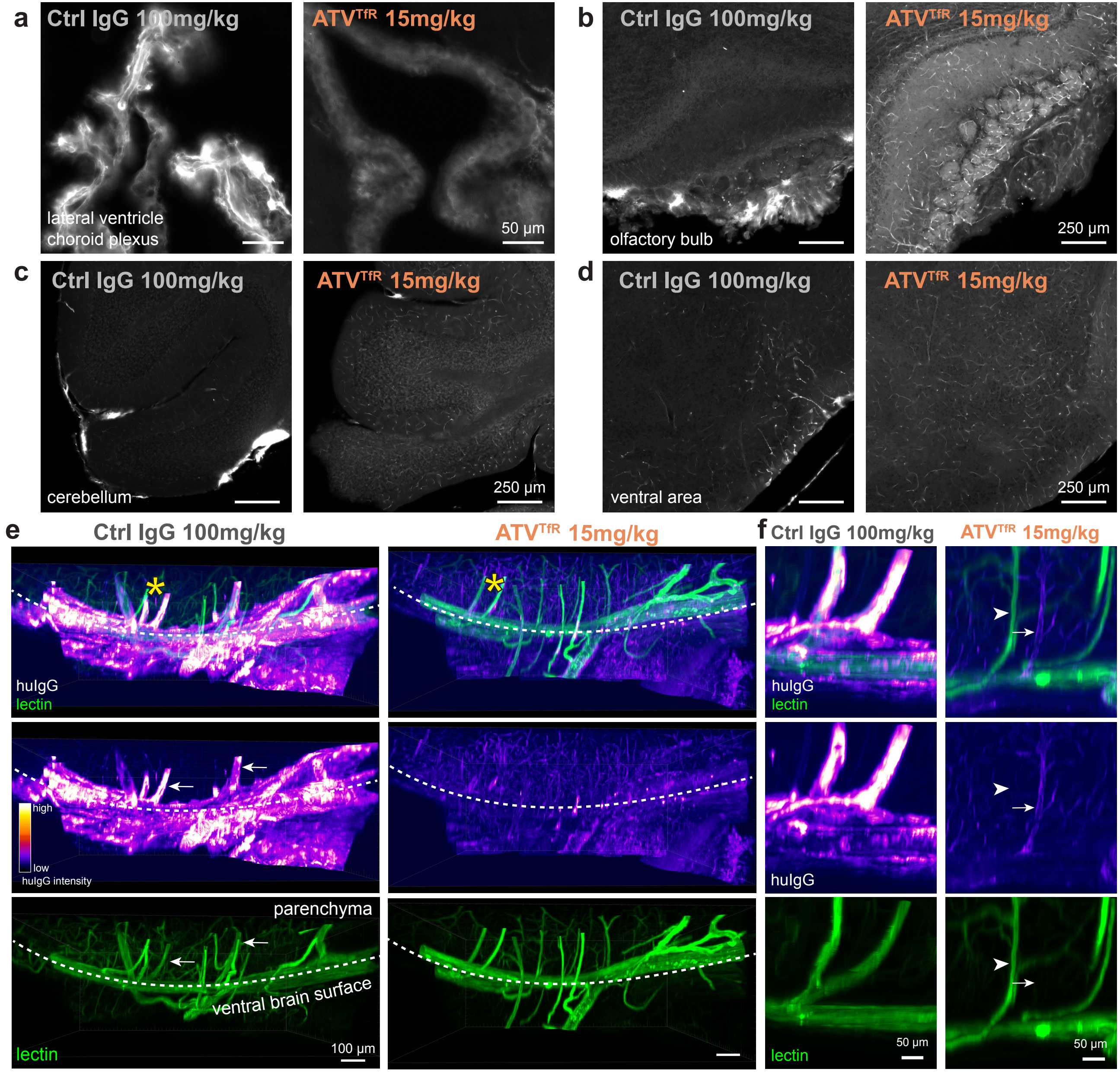

**Supplementary Figure 1. Increasing control IgG dose to match ATV$^{TfR}$ brain concentration does not improve its limited biodistribution**. **a-d.** Representative Immunofluorescence images of administered huIgG in brain-concentration-matched mouse study in indicated brain regions (see **Fig. 3d-h**, n=6/group). Brightness settings were adjusted due to extremely high signal in the choroid plexus for control IgG. **e.** Lateral 3D view of the circle of Willis and penetrating vessels; arrows indicate perivascular signal around penetrating vessels and asterisks indicate approximate vessels captured in **Fig. 3f**. **f**. Optical slice (200 μm) of penetrating parenchyma vessels; lectin-positive putative arterioles (arrowhead) and lectin-negative, ATV-positive putative venule (arrow). For micrographs, display settings were optimized independently for each region of interest and were kept identical for both treatment groups.

Supplementary Figure 2

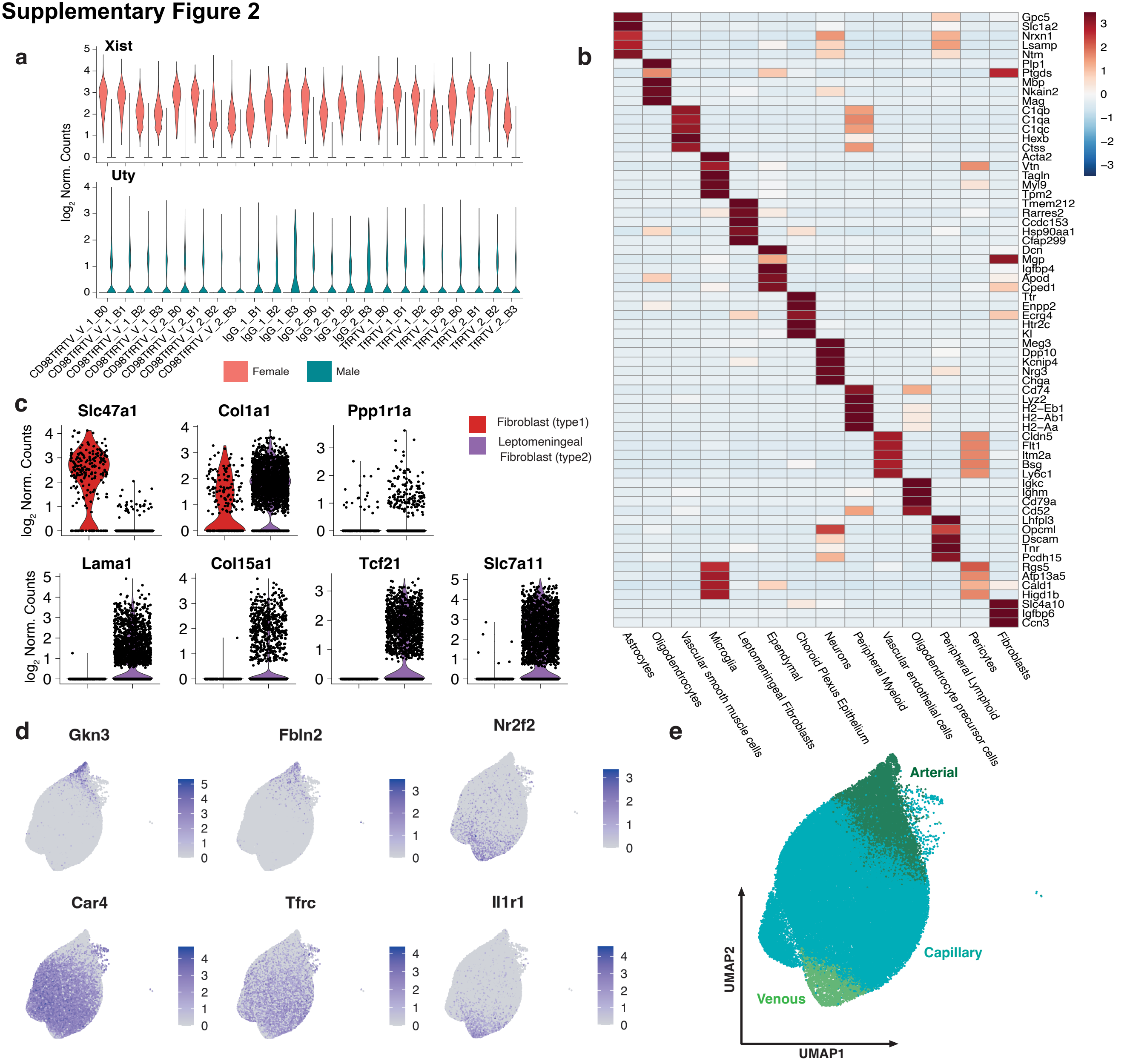

**Supplementary Figure 2. Expression levels of markers used for scRNAseq cellular annotations**
**a.** Violin plots of the distribution of X chromosome gene *Xist* and Y chromosome gene *Uty* of each individual animal from experiment in Fig. 4-5. Distributions support successful separation of individual mice based on predicted sex. **b.** Heatmap showing average expression of top 3 marker genes for each predicted class of cells. **c.** Violin plots of the distribution of established markers of type 1 and type 2 fibroblasts. **d.** UMAP plots colored according to log normalized expression for established markers of arterial-capillary-venous axis. **e.** UMAP showing arterial-capillary-venous axis in vascular endothelial cells.

**Supplementary Figure 3**

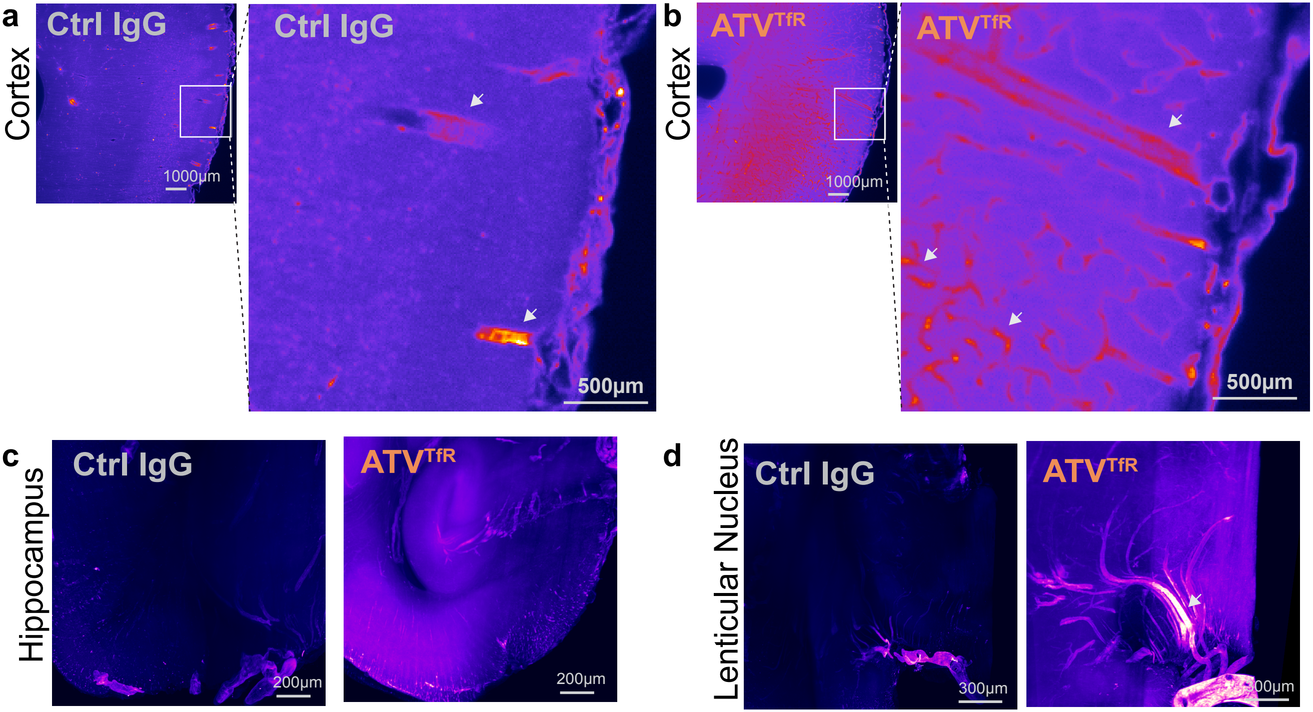

**Supplementary Figure 3**. **Enhanced exposure and biodistribution of ATV$^{TfR}$ in the cortex, hippocampus, and lenticular nucleus of cynomolgus monkey brains. a-d.** Representative coronal slice view (200 µm thick) immunofluorescence images of cortex (a-b), hippocampus (c), and lenticular nucleus (d) from cynomolgus monkeys dosed with either AF647-conjugated control IgG **(a)** or ATV$^{TfR}$ **(b)** 2 days post-dose. Arrows in (a) highlight localization of control IgG mostly to large penetrating vessels while arrows in (b) highlight uptake of ATV$^{TfR}$ within both large vessels as well as smaller capillaries. Arrow heads in (d) highlighting the localization of ATV$^{TfR}$ to the habenular nucleus. n = 1/group.

#### Supplementary Figure 4

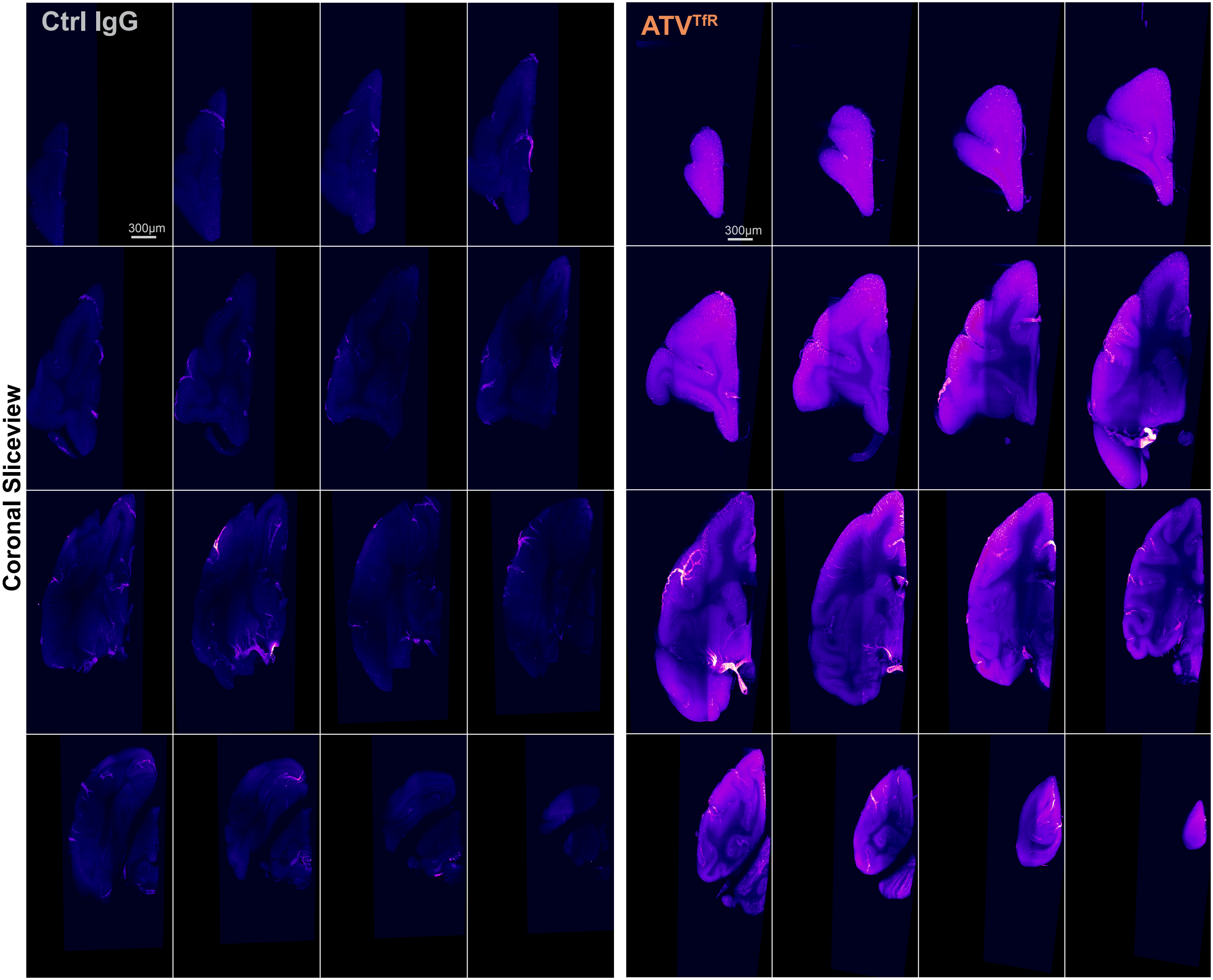

**Supplementary Figure 4**. **Enhanced exposure and biodistribution of ATV$^{TfR}$ in the brain cynomolgus monkey across multiple brain regions.** Representative coronal slice view (200 µm thick) across cynomolgus monkey brain from rostral to caudal from cynomolgus monkeys dosed with either AF647-conjugated control IgG or ATV$^{TfR}$ 2 days post-dose. n = 1/group.
