## Supplementary Table 1 for "Fc-engineered large molecules targeting blood-brain barrier transferrin receptor and CD98hc have distinct central nervous system and peripheral biodistribution compared to standard antibodies"

| **Region of Interest** | **CCFv3 IDs** | **CCFv3 Names** |
| --- | --- | --- |
| olfactory bulb | 507, 151 | main olfactory bulb, accessory olfactory bulb |
| anterior olfactory nucleus | 159 | anterior olfactory nucleus |
| cortex | 315 | isocortex |
| hippocampus | 1080 | hippocampal region |
| septum | 250, 258, 581, 310, 564 | lateral septal nucleus, caudal (caudodorsal) part, lateral septal nucleus, rostral (rostroventral) part, triangular nucleus of septum, septofimbrial nucleus, medial septal nucleus |
| caudoputamen | 672 | caudoputamen |
| ventral striatum | 465, 458, 473, 56, 342, 298, 596, 481 | olfactory tubercle, pyramidal layer, olfactory tubercle, molecular layer, olfactory tubercle, polymorph layer, nucleus accumbens, substantia innominata, magnocellular nucleus, diagonal band nucleus, islands of calleja |
| thalamus | 549 | thalamus |
| hypothalamus | 1097 | hypothalamus |
| midbrain | 313 | midbrain |
| pons | 771 | pons |
| medulla | 354 | medulla |
| cerebellum | 512 | cerebellum |
| bed nuclei of the stria terminalis | 351 | bed nuclei of the stria terminalis |
